# Next-gen glycoengineering: combining cellular and metabolic engineering to fine-tune mAb β1,4-galactosylation

**DOI:** 10.64898/2026.09.26.754714

**Authors:** Apostolos Tsopanoglou, Mina Ghahremanzamaneh, Itzcóatl Gómez Aquino, Kallum Doyle, Brian Glennon, Sara Carillo, Jonathan Bones, Ioscani Jiménez del Val

## Abstract

N-linked Fc galactosylation drives variability in therapeutic mAb quality, affecting complement-dependent cytotoxicity and ADCC. Current control strategies suffer from narrow ranges, productivity loss, or unwanted glycoforms. Here, we combine metabolic and cellular glycoengineering by feeding 2-deoxy-2-fluorogalactose (2FG) to CHO cells engineered for hypergalactosylation via COSMC knockout and β4-galactosyltransferase overexpression (C-/GT+), achieving broad, tuneable control of Fc β4-galactosylation. In DP12 C-/GT+ cells, a split-feed regimen spanned a wide galactosylation range (95-46%) but generated substantial aglycosylated and Man5 glycoforms. The same regimen in a second host, VRC01 C-/GT+, gave only a narrow range, indicating 2FG dosing does not transfer directly across cell lines. Based on a proposed mechanism where 2FG is metabolised via the Leloir pathway into inhibitory fluorinated nucleotide-sugar intermediates, we developed a cell-normalised, multi-bolus feeding strategy that substantially broadened control in VRC01 C-/GT+ cells while suppressing aglycosylation and Man5 formation. In a fed-batch Ambr^®^ 250 process, this strategy achieved comparable galactosylation reduction with roughly ten-fold less 2FG, showing that the fraction of mAb secreted after feeding is a key determinant of control strategy efficiency. Across the resulting glycoform range, Bio-Layer Interferometry identified Fc β4-galactosylation as the strongest correlate of FcγRIIIA (CD16a) binding affinity (R^2^ = 0.92). Together, these findings establish combined metabolic and cellular glycoengineering as a platform for controlling mAb galactosylation across a broad range, with potential to fine-tune downstream pharmacological activity.

**Highlights:**

- Cell-normalised 2FG feeding tunes mAb galactosylation
- C-/GT+ hypergalactosylation extends the achievable control range
- Post-feed production timing governs 2FG feeding efficiency
- Fed-batch bioreactors need ∼10-fold less 2FG for an equivalent control range
- Galactosylation is the strongest correlate of CD16a binding affinity

## 1. Introduction

Monoclonal antibodies (mAbs) have had annual sales exceeding US $200 billion since 2021 (Grand View Research, 2026; Walsh and Walsh, 2022) and continue to have a strong growth forecast over the next decade (Crescioli et al., 2026; Grand View Research, 2026). Over 200 approved mAbs continue to address critical diseases such as cancer, immune disorders, infectious diseases, and genetic disorders (Crescioli et al., 2026; Walsh and Walsh, 2022). With an annual rate of over 18 mAb products gaining first marketing approval since 2023, their economic significance and life-saving potential continue expanding (Crescioli et al., 2026).

The average commercial-scale mammalian cell culture titre is now above 5 g/L in extended fed-batch cultures, increasing significantly over the past decade (Shukla et al., 2017). This indicates that productivity, once a significant concern, has become less of a barrier. Instead, product quality assurance has become the key driver for this multibillion-dollar industry, motivating ongoing efforts to minimise mAb product variability.

N-linked β4-galactosylation is a major source of heterogeneity in commercial mAbs, varying across different lots of blockbuster products (Planinc et al., 2017). mAbs contain variable levels of non- (A2G0) or mono-galactosylation (A2G1) (Luo et al., 2026), and increased Fc galactosylation has been shown to enhance complement-dependent cytotoxicity (CDC) (van Osch et al., 2021) antibody-dependent cellular cytotoxicity (ADCC) (Thomann et al., 2016), and antibody-dependent cellular phagocytosis (ADCP) (Kuhns et al., 2020), all important effector functions that underlie the oncolytic activity of many therapeutic mAbs. Despite the clear need to tightly control mAb Fc β4-galactosylation and ensure a consistent therapeutic profile, strategies to achieve this remain limited.

Cell engineering strategies have been developed to ablate or enhance mAb Fc β4-galactosylation through knockout or overexpression of single or multiple genes of the β1,4-galactosyltransferase glycoenzyme family (Amann et al., 2018; Nguyen et al., 2021; Raymond et al., 2015). However, such approaches lead to set galactosylation levels that are not amenable to real-time control applications. More recently, advanced cell engineering strategies have emerged, where inducible transcriptional circuits have been used to regulate expression of glycosylation enzymes via addition of small molecule inducers to culture (Chang et al., 2019).

Metabolic glycoengineering strategies to modulate mAb β4-galactosylation have also been developed. In these, uridine and galactose, which are direct metabolic precursors of uridine diphosphate galactose (UDP-Gal), the obligate co-substrate for galactosylation reactions, are fed alongside manganese, the cognate co-factor of β4-galactosyltransferases, to drive mAb Fc β4-galactosylation (Grainger and James, 2013; Gramer et al., 2011). However, the UMG feeding strategy only achieves a maximum control range of (∼25%) at the expense of reduced cell growth or product yield and, in some instances, increased production of undesired high mannose glycans (Grainger and James, 2013; Pranomphon et al., 2025).

An alternative strategy that has been explored involves feeding dehydroxylated or fluorinated galactose analogues to inhibit galactosylation. Originally, galactose analogues were used to fully ablate glycosylation in rat hepatocytes and evaluate the impact of truncated glycans on cell membrane integrity and the properties of secreted proteins (Gross et al., 1992; Loch et al., 1991). Dekkers et al. (2016) revisited this strategy by feeding 2-deoxy-2-fluoro-D-galactose (2FG) to mAb-producing HEK293 cells, achieving dose-dependent control of β4-galactosylation from 9% to 26%, with the upper value corresponding to unfed wild-type levels. Inspired by the seminal work of Dekkers et al. (2016), the present study extends 2FG feeding to CHO cells engineered to yield mAb Fc hypergalactosylation via combined knockout of the core 1 β3-galactosyltransferase-specific molecular chaperone (C-) and β4-galactosyltransferase overexpression (GT+) (Gomez Aquino et al., 2027), with the aim of achieving a broader range of β4-galactosylation control than is attainable in wild-type cells. Initial evaluation in DP12 C-/GT+ cells achieved a broad galactosylation control range (95% down to 46%) but at the expense of substantial aglycosylated and Man5 mAb Fc production, while the same feeding regime applied to a second hypergalactosylating cell line, VRC01 C-/GT+, yielded only a narrow control range, revealing that a fixed, volume-normalised 2FG dose is not directly transferable across cell lines.

Guided by a proposed mechanism whereby 2FG perturbs the Leloir pathway and nucleotide sugar metabolism, a cell-normalised, multi-bolus feeding strategy was developed that substantially broadened the achievable control range in VRC01 C-/GT+ cells (95% down to 57%) while alleviating off-target glycoform production. This same strategy, deployed in Ambr^®^ 250 bioreactors operated in fed-batch mode, achieved a comparable galactosylation reduction at roughly an order of magnitude lower 2FG exposure, highlighting production-phase dynamics as a critical, previously unrecognised variable governing feeding efficiency. Samples spanning the obtained glycoform range demonstrated that FcγRIIIA (CD16a) binding affinity, a key determinant of ADCC activity, correlates most strongly with Fc β4-galactosylation.

Collectively, these findings establish the combination of C-/GT+ glycoengineering and optimised 2FG feeding as a robust strategy to profile the pharmacological impact of defined galactosylation setpoints, with potential applications in novel mAb development and biosimilar design.

## 2. Material and Methods

### 2.1. Reagents

2-deoxy-2-fluoro-D-galactose (2FG) was purchased from Carbosynth (Cat. No. MD04718). A working stock of 68.62 mM was prepared by dissolving 2FG in water followed by filtering with a 0.22 μm Millipore filter. 2FG was fed into the culture at the specified time points as highlighted in the main text.

### 2.2. CHO cell culture

To evaluate our proposed strategy in this study two mAb-producing CHO cell lines were used. CHO DP12 clone #1934 (ATCC^®^ CRL-12445™) producing an anti-interleukin-8 mAb (Heinrich et al., 2011), which was purchased from LGC Standards (Middlesex, UK) and adapted to serum-free suspension culture at the National Institute for Bioprocessing Research and Training (NIBRT), was grown in Ex-Cell^®^ 302 (Sigma-Aldrich, Cat. No. 14324C) medium, supplemented with 4 mM glutamine (Merck Millipore, Cat. No. K0283-BC) and 200 nM methotrexate (Sigma-Aldrich, Cat. No. 454126). The second cell line, CHO VRC01, expresses a broadly neutralising antibody (bNAb) targeting HIV-1 CD4 binding sites (Wu et al., 2010) and was generously gifted by the US National Institutes of Health. VRC01 cells were grown in ActiPro^TM^ medium (Cytiva/HyClone, Cat. No. SH31039.02) supplemented with 4 mM of L-glutamine and 100 nM methotrexate. The anti-interleukin-8 mAb has a conserved glycosylation site on Asn297 of the Fc while, interestingly, the bNAb contains an additional glycosylation site in its Fab fragment.

During routine passaging, cells were seeded at 0.2 million cells/mL every 2 days and propagated to a VCD of 1.2-1.5 million cells/mL in T-25 non-surface treated flasks (Fisher Scientific, Cat. No. 10476921). If not stated otherwise, suspension cultures were conducted in 125 mL vented shake flasks (Fisher Scientific, Cat. No. 10266432) with a 30 mL working volume and a seeding density of 0.2 million cells/mL on a shaking platform rotating at 120 rpm with a 9.5 mm orbital radius (Thermo Fisher Scientific, Waltham, MA, US, Cat. No. 88881102), at 5% CO_2_ saturation and 37 °C in a Steri-Cycle CO_2_ incubator (Thermo Fisher Scientific, Waltham, MA, US). Cultures were terminated when viability fell below 70%. Viable cell density (VCD) was determined through dye exclusion with a haemocytometer and 0.4% (w/v) trypan blue (Gibco, Cat. No. 15250061).

### 2.3. Ambr^®^ 250 bioreactor runs

Bioreactor runs were performed in Ambr^®^ 250 bioreactors (Sartorius Stedim, Göttingen, DE) with two pitched blade impellers and an open pipe sparger (Sartorius, Cat. No. 001-5G25). Bioreactor inoculation was at 0.4 million cells/mL in a working volume of 210 mL of BalanCD CHO Growth A media (Irvine Scientific, Cat. No. 91128). BalanCD CHO Feed 4 (Irvine Scientific, Cat. No. 94134) was added on days 3, 5, 7, and 9 (6% v/v of the original working volume) as per the manufacturer’s instructions. In addition, a 1 M glucose solution was fed, as required, to maintain glucose above 4 g/L. Temperature was set at 36.5 °C, and a proportional-integral-derivative (PID) controller was used to maintain dissolved oxygen (DO) at 50% of air saturation. Impeller stirring speed was set to 400 rpm. pH was set to 7.0 ± 0.1, with base addition (1 M sodium carbonate) controlled through proportional control to a deadband of 0.1 pH units, and CO_2_ sparging controlled under Proportional-Integral (PI) cascade control, with integral control in place of a specified deadband. A Vi-CELL XR (Beckman Coulter, Brea, CA, US) for measuring viable cell density, and a BioProfile^®^ FLEX2 cell culture analyser (Nova Biomedical, Waltham, MA, US) for analysing cell culture parameters (glucose, lactate, glutamine, glutamate, NH_4_^+^, pH, pCO_2_, and pO_2_) were integrated with the Ambr^®^ 250 system.

### 2.4. Antibody quantification

Antibody titre was quantified from cell culture supernatant, using an Agilent 1200 series High-Performance Liquid Chromatography (HPLC) instrument (Agilent Technologies, Santa Clara, CA, US) with a UV detector and a Thermo Scientific MAbPac™ Protein A column (Thermo Scientific, Cat. No. 082539). A pH gradient program was applied for elution, where eluent A was 50 mM sodium phosphate and 150 mM sodium chloride pH 7.5 and eluent B was 50 mM sodium phosphate and 150 mM sodium chloride pH 2.5. The following elution gradient was used: t_0-3(min)_ = 0% B, t_3-5(min)_ = 100% B, t_5-8(min)_ = 0% B. Detection was performed at 280 nm. Prior to injection, 0.5 mL of culture broth was collected, spun down at 3,000 ×g for 10 minutes and 0.2 μm filtered (Fisher Scientific, Cat. No. 15342378).

### 2.5. Antibody purification and glycan analysis

On the final day of culture, ∼15 mL were harvested by spinning down the broth at 3,000 ×g for 10 minutes. The supernatant was processed on an ÄKTA Start^TM^ FPLC (Cytiva, Uppsala, SE) fitted with a HiTrap^®^ protein A column (Cytiva, Cat. No. 17040201). After elution, pH was neutralised with 200 μL of 1M Tris-HCL (pH 9) and concentration was determined with a ND-100 Nanodrop^TM^ spectrophotometer (Thermo Fisher Scientific, Paisley, UK).

Single-chain Fc (scFc) analysis workflow was adapted from the method previously developed by Carillo et al. (2020). Briefly, mAb subunits, generated by digestion with the immunoglobulin-degrading enzyme of *Streptococcus pyogenes* (IdeS) (Genovis, Kävlinge, Sweden), were separated on a Vanquish^TM^ Flex Binary UHPLC instrument (Thermo Fisher Scientific, Sunnyvale, CA, US) equipped with a MAbPac^TM^ RP column (2.1 × 50 mm, 4 µm; Thermo Fisher Scientific, Cat. No. 088648) and coupled to a Thermo Scientific Orbitrap Exploris 240 mass spectrometer (Thermo Fisher Scientific, Bremen, DE) for high-resolution accurate-mass analysis of Fc subunits. Raw data were deconvoluted using Xtract algorithm in BioPharma Finder^TM^ software (Thermo Fisher Scientific, San Jose, CA, US). Further details on LC-MS set up and data analysis are provided in the Supplementary Section 1.

### 2.6. Bio-layer Interferometry

Bio-Layer Interferometry (BLI) binding assays were performed by Sino Biological (Beijing, China) to determine the affinity of selected mAb glycoform samples for recombinant human CD16a (FcγRIIIA), using an Octet^®^ RED384 system (ForteBio/Sartorius, Fremont, CA, US). Recombinant CD16a was immobilised onto Anti-Penta-HIS (HIS1K) biosensors (Sartorius, Cat. No. 18-5120) at 2 µg/mL until a loading response of approximately 1 nm was reached. Loaded sensors were dipped into serial dilutions of each mAb sample (6.25 to 100 nM) for association (240 s), followed by dissociation in kinetics buffer (300 s), at 20°C with agitation at 1000 rpm. A buffer-only reference sensor was included to correct for baseline drift and non-specific binding. Data were double-referenced and fitted to a 1:1 binding model using the Octet^®^ Data Analysis software (version 12.0.1.2) to determine *k_on_*, *k_off_*, and equilibrium dissociation constants (*K_D_*). Duplicate measurements were performed on each sample.

### 2.7. Statistical analysis & calculations

Statistical analysis was performed using GraphPad Prism 11.0.2. The level of significance was set at p < 0.05 using ANOVA followed by Tukey’s test for pairwise mean comparisons. *, **, ***, and **** denote p ≤ 0.05, p ≤ 0.01, p ≤ 0.001, and p ≤ 0.0001, respectively.

The fraction of mAb Fc glycoform *i* secreted after 2FG feeding, *f_i_*, was calculated with Equation 1 (Fan et al., 2015), where *x_i_* is the measured fraction of glycan *i* at the culture endpoint, *x_i_*_,*UF*_ is fraction of glycan *i* at the endpoint of the corresponding (unfed) control culture, [*mAb*]*_f_* is the molar concentration of mAb at the end of culture, and [*mAb*]_72_is the mAb molar concentration at 72 hours of culture (immediately before the first 2FG feed). Equation 1 assumes that the glycosylation profile secreted by the cells before the first 2FG is the same as the one produced over the full culture by unfed cells.

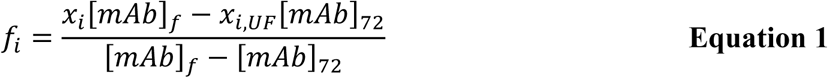

## 3. Results and Discussion

### 3.1. Initial evaluation of 2FG feeding of C-/GT+ CHO cells

The C-/GT+ CHO cell lines contain a dual genetic modification designed to achieve hypergalactosylation of mAb Fc N-glycans (Gomez Aquino et al., 2027). This approach couples the knockout of COSMC, the molecular chaperone required for functional C1GalT1 activity, with ectopic expression of human β4GalT1. Loss of COSMC abolishes O-linked β3-galactosylation, thereby redirecting the cellular UDP-Gal pool towards N-linked β4-galactosylation of the mAb Fc, while β4GalT1 overexpression alleviates the enzymatic bottleneck that would otherwise limit galactosylation capacity. Together, these two engineering events act synergistically, one increasing UDP-Gal donor availability for Fc glycosylation and the other increasing acceptor enzyme availability, to yield >90% Fc β4-galactosylation in both DP12 and VRC01 CHO cell lines (Gomez Aquino et al., 2027). Given this hypergalactosylation phenotype, the C-/GT+ glycoengineering provides an ideal platform to investigate strategies that modulate mAb galactosylation. In particular, this system was used to evaluate feeding of 2-deoxy-2-fluorogalactose (2FG) as an actuator to titrate mAb Fc β4-galactosylation from high to low levels.

Initial experiments with the DP12 C-/GT+ cell line focussed on evaluating the impact of 2FG feeding on cell growth, given that Leloir pathway inhibition can potentially perturb general metabolic flux beyond its intended effect on Fc galactosylation. Three feeding regimes were compared: (i) a full 2FG dose administered on day 0 of culture, (ii) a full 2FG dose administered on day 3 of culture, and (iii) a split-dose regimen consisting of 10% (w/w) of the total dose on day 3 followed by the remaining 90% (w/w) on day 5, as previously described by Dekkers et al. (2016). This comparison allowed both the timing and the dosing schedule of 2FG addition to be assessed for their respective effects on culture viability and growth kinetics, providing a basis for selecting a feeding strategy that minimises any adverse impact on cell growth while retaining the capacity to modulate mAb Fc β4-galactosylation.

Supplementary Figure 1 presents VCD for the three feeding regimes. Day 0 2FG feeding severely impacted VCD across all doses. Full 2FG dosing on day 3 also reduced VCD at 2FG concentrations beyond 0.1 mM, and the 10:90 split-feed regime had no observable impact on VCD up to 0.5 mM 2FG. Inhibition of cell proliferation with 2FG feeding during exponential growth is likely due to disruption of glycolysis. CHO cells present high glycolytic flux during exponential growth (Ahn and Antoniewicz, 2011), and this pathway provides ATP at a rate that matches demand for high proliferation (Kukurugya et al., 2024).

Simultaneously, CHO cells achieve comparable growth when galactose replaces glucose as the main carbon source (Torres et al., 2019). It follows that, like galactose, 2FG can be efficiently channelled through the Leloir pathway to yield 2-deoxy-2-fluoro glucose-6-phosphate (2FGlc-6P). Because 2FGlc-6P lacks the 2-hydroxyl group required for conversion to fructose-6-phosphate, it accumulates within the cells (Gallagher et al., 1978) and may stall glycolytic activity and, thereby, proliferation. This would explain why the split-feed regime, which delays the bulk of the dose until later in culture, showed no impact on VCD and was consistent with Dekkers et al. (Dekkers et al., 2016). This mechanism remains speculative and would require validation through intracellular metabolite analysis (e.g., LC-MS/MS or 19F-NMR) and/or direct assays of glycolytic enzyme activity in 2FG-treated cells.

### 3.2. 2FG feeding modulates mAb Fc glycosylation in DP12 C-/GT+ cells

Once a feeding schedule that avoided reduced cell proliferation was established, a wide range of 2FG concentrations (0.1 mM – 2 mM) was evaluated around key cell culture indicators and, most importantly, the decoy substrate effect on final mAb product glycosylation.

The assessed 2FG concentrations had no discernible effect on cell growth (Figure 1A), and small improvements in specific productivity (q_p_) were observed with increasing 2FG dosing (table inlay in Figure 1B). Combined with constant VCDs, the higher q_p_ translated into higher product titre at the upper 2FG dose range (≥1.0 mM) (Figure 1B). Flow cytometry analysis revealed increased cell size in the high 2FG concentration conditions (data not shown). Larger cell volume has been linked with increased specific productivity in CHO cells (Pan et al., 2017).

**Figure 1.**
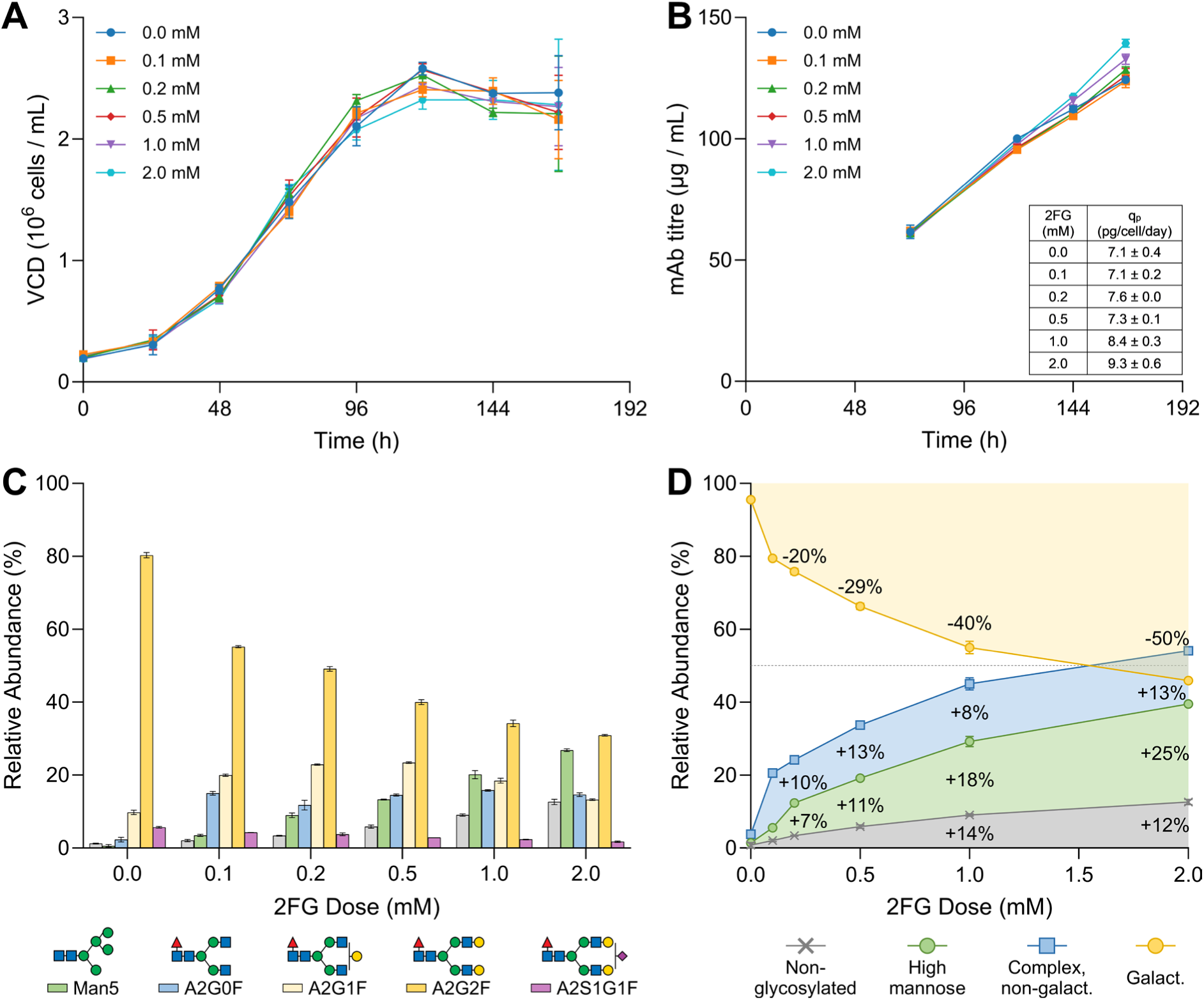
Culture performance of DP12 C-/GT+ cells fed with varying doses of 2FG. Viable cell density (A), mAb titre (B), glycan profile (C), and glycan motifs (D) are presented for CHO DP12 C-/GT+ cells treated with the 10:90 split feed (0.0–2.0 mM 2FG). Data are mean values ± standard deviation for biological duplicates (N = 2). The glycan notation used in (C) corresponds to the Consortium for Functional Glycomics (CFG) nomenclature (Varki et al., 2015). In (D), yellow corresponds to the sum of galactosylated species (A2G1F + A2G2F + A2S1G1F), blue is the A2G0F complex, non-galactosylated glycan, green correspond to the Man5 glycan, and grey is aglycosylated mAb Fc. The negative quantities shown in (D) correspond to reductions in mAb Fc galactosylation with increasing 2FG doses, while the positive values indicate the concomitant increase in non-galactosylated glycan motifs.

mAb Fc glycoprofiling revealed a dose dependent decrease in galactosylated A2G1F and A2G2F species, along with a concomitant increase in Man5 and A2G0F glycoforms (Figure 1C). The fraction of total mAb Fc galactosylated glycoforms decreased with increasing 2FG dose from 95.5% ± 0.2% down to 45.9% ± 0.2%, a difference of 49.6% ± 0.2% (Figure 1D). Figure 1D also shows the contribution of different glycan classes to the galactosylation reductions. As 2FG doses increase, aglycosylation and Man5 glycans are the major contributors to the observed reductions in galactosylation, while non-galactosylated complex glycans remain relatively consistent across 2FG doses, increasing between 8% and 13% across the 2FG dose range. The considerable increases in aglycosylated and Man5 glycans were unanticipated, given that they were not observed in previous work with 2FG (Dekkers et al., 2016).

### 3.3. 2FG feeding modulates mAb Fc glycosylation in VRC01 C-/GT+ cells

To determine whether the effect of 2FG feeding on VCD, titre, and mAb Fc glycosylation is cell-line dependent, the same feeding regime (10%:90% split across days 3 and 5) and 2FG dose range were tested on a second hypergalactosylating cell line, CHO VRC01 C-/GT+.

In VRC01 C-/GT+ cells, the 2FG feeding regime and concentration range yielded only minor differences in VCD, with slightly lower values at the endpoint of culture at higher 2FG doses (Figure 2A). No differences were observed in mAb titre or q_p_ (Figure 2B). Interestingly, the impact of 2FG doses on glycosylation were reduced substantially in VRC01 C-/GT+ cells, with only minor reductions in A2G1F and A2G2F species with 2FG doses up to 0.2 mM (Figure 2C). At 2FG doses ≥ 0.5 mM, larger reductions in A2G2F were obtained. The maximum reduction in overall mAb galactosylation under these conditions was 22% and arose, largely due to an increase in A2G0F (17%) and only a small contribution from Man5 (4%) (Figure 2D). Interestingly, production of aglycosylated mAb Fc was not detected.

**Figure 2.**
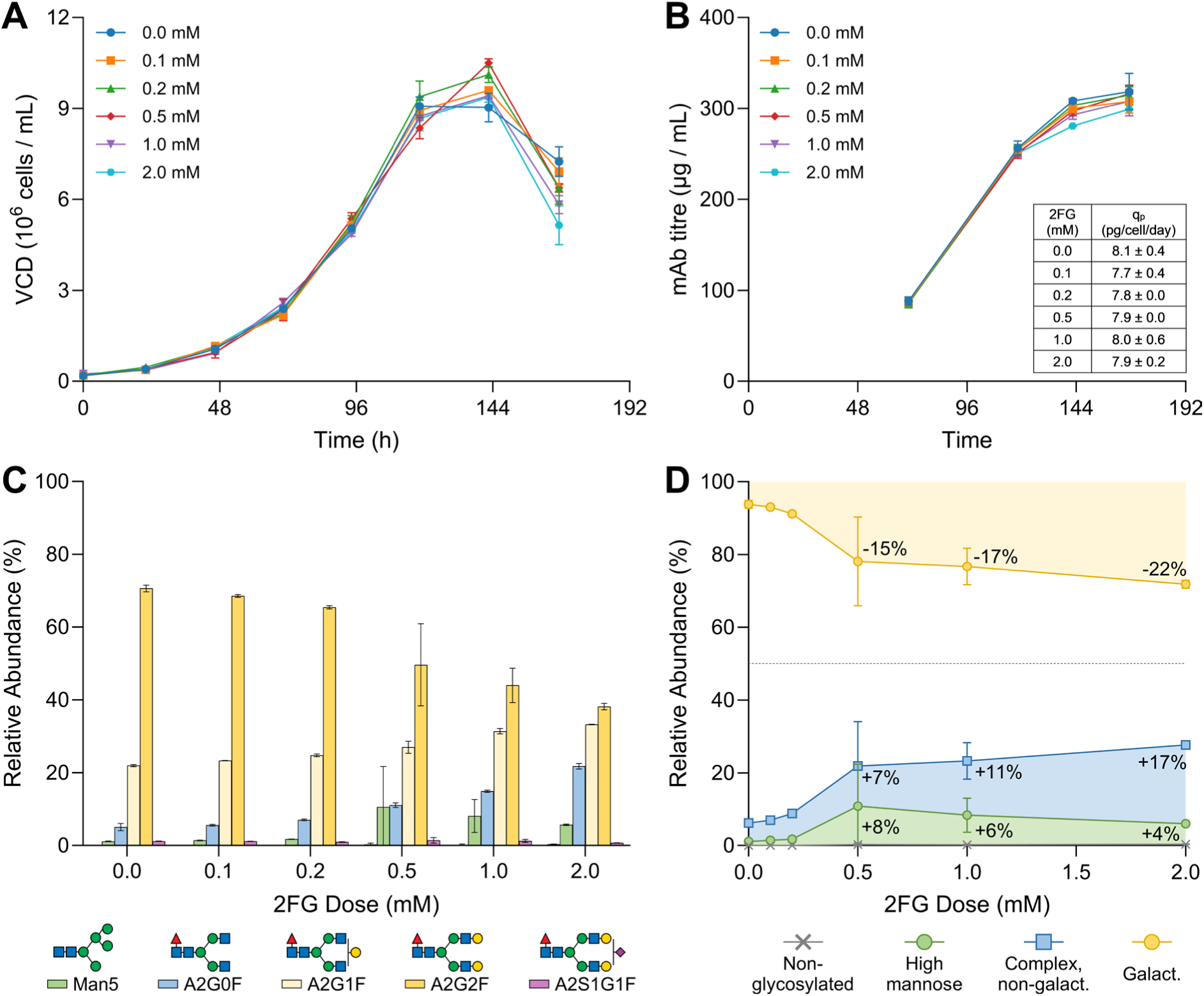
Culture performance of VRC01 C-/GT+ cells fed with varying doses of 2FG. Viable cell density (A), mAb titre (B), Fc glycan profile (C), and Fc glycan motifs (D) are shown for CHO VRC01 C-/GT+ cells treated with the 10:90 split feed of 2FG (0.0-2.0 mM). Values represent means ± standard deviation of biological replicates (N = 2). Glycans in (C) follow CFG notation (Varki et al., 2015). In (D), yellow denotes the sum of galactosylated glycans, blue the complex, non-galactosylated A2G0F glycan, green Man5, and grey aglycosylated mAb Fc. Negative values reflect the loss of Fc galactosylation while positive values reflect the corresponding rise in non-galactosylated motifs.

### 3.4. Enhancing 2FG feeding

In DP12 C-/GT+ cells, 2FG feeding leads to high production of aglycosylated and Man5 mAb Fc glycoforms. Aglycosylated mAbs present depleted ADCC and CDC activity through impaired FcγR binding (Tao and Morrison, 1989). Man5 mAb glycoforms present reduced serum half-life through clearance via mannose receptors (Baumeister et al., 2026). Man5 glycans also raise immunogenicity concerns through the possibility of lectin-mediated immune recognition (Dasgupta et al., 2007; Delignat et al., 2020). Moreover, the 2FG dosing that greatly modulated glycosylation in DP12 cells had a considerably lower impact in VRC01 C-/GT+ cells. Both the production of aglycosylated and Man5 glycans along with the disparate impact on glycosylation across cell lines highlighted the opportunity to optimise 2FG feeding to (i) minimise production of deleterious mAb glycoforms and (ii) define a strategy to achieve similar glycosylation tuning across different cell lines.

To optimise 2FG feeding, the mechanisms by which it inhibits galactosylation and causes aglycosylation and Man5 production must be defined. Figure 3 outlines the proposed mechanisms, contrasting baseline galactose metabolism with 2FG feeding, based on the substrate specificity of Leloir pathway enzymes (Holden et al., 2003), structural analogy between fluorinated and non-fluorinated carbohydrates, and the nucleotide sugar donor biosynthetic pathway (Freeze et al., 2022). A recent HEK293 cell study found that fluorinated galactose 1-phosphate (dGalF-1P) competes with galactose 1-phosphate for the binding sites of GALT (Janes et al., 2020), suggesting this enzyme as the point where 2FG enters the Leloir pathway (Figure 3).

**Figure 3.**
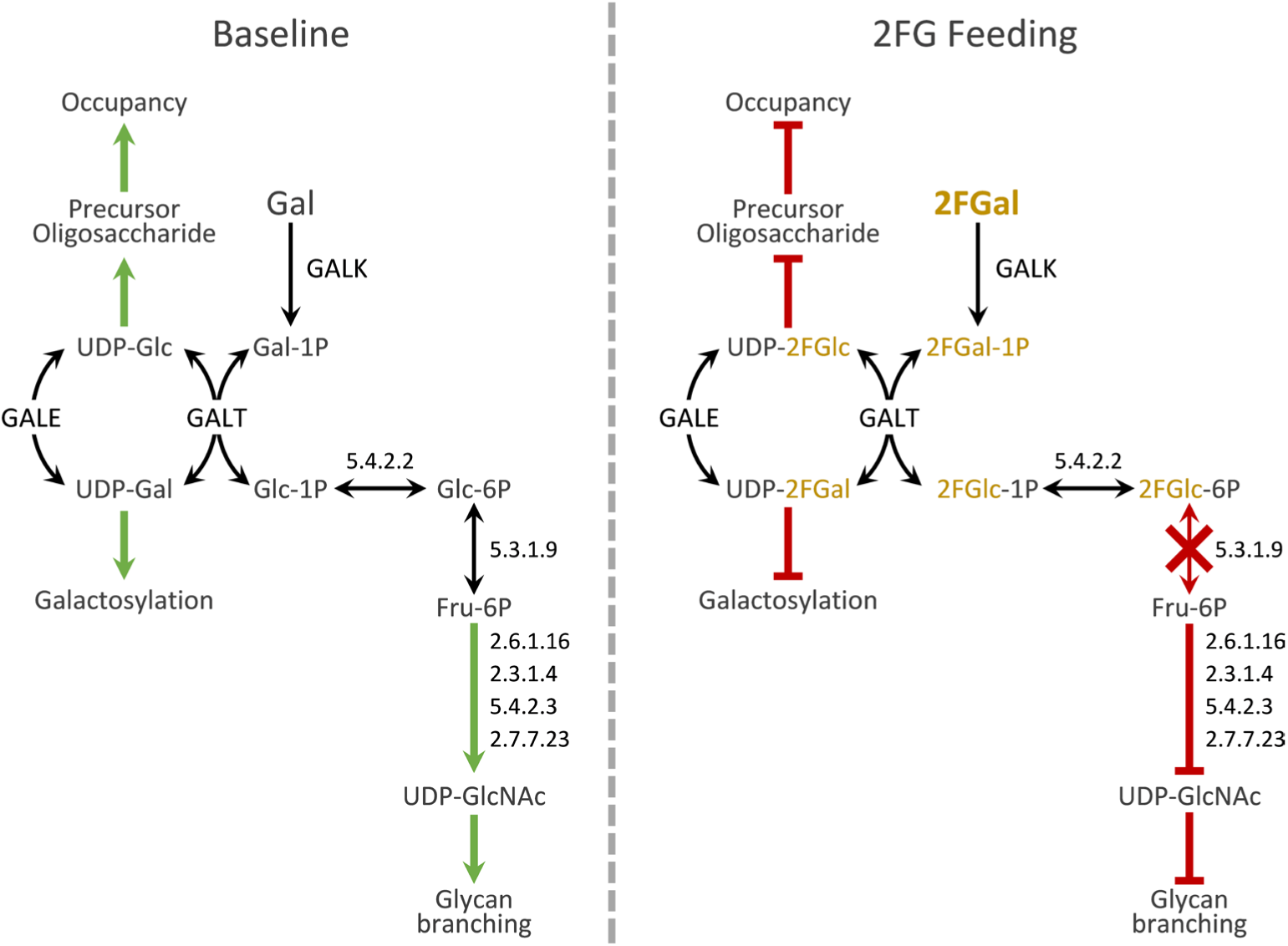
Proposed mechanism of 2FG feeding on CHO cell glycosylation. Under baseline conditions, galactose that is fed to culture is phosphorylated by GALK to produce Gal-1P, which enters the Leloir pathway. GALT exchanges the UDP nucleotide with UDP-Glc to yield Glc-1P and UDP-Gal. UDP-Gal can then serve as a co-substrate for galactosylation reactions within Golgi (Freeze et al., 2022). UDP-Gal can also be epimerised by GALE to yield UDP-Glc. In turn, UDP-Glc serves as a co-substrate to produce the Glc_3_Man_9_GlcNAc_2_ precursor oligosaccharide that is transferred to a nascent polypeptide chain in the first step of the N-linked glycosylation process (Stanley et al., 2022). Glc-1P, produced from the first step of the Leloir pathway (Holden et al., 2003) can then be epimerised to Glc-6P and then Fru-6P which can yield pyruvate, which feeds into the TCA cycle (Ahn and Antoniewicz, 2011). Fru-6P also serves as the main carbon backbone substrate for UDP-GlcNAc synthesis (Freeze et al., 2022), which is the nucleotide sugar that underlies N-glycan branching (Stanley et al., 2022). Although not experimentally confirmed, we propose, based on the known substrate specificity of Leloir pathway enzymes and the structural similarity between 2FG and its non-fluorinated counterpart, that when 2FG is fed to culture, it is phosphorylated by GALK and enters the Leloir pathway to yield UDP-2FGal, which inhibits protein galactosylation. UDP-2FGal can then be epimerised by GALE to yield UDP-2FGlc, which may inhibit synthesis of the Glc_3_Man_9_GlcNAc_2_ precursor oligosaccharide and, thereby, reduce N-glycan occupancy (Liu et al., 2014). UDP-2FGlc can continue through the Leloir pathway to yield 2FGlc-1P and, subsequently, 2FGlc-6P. Absence of the hydroxyl group at C2 in 2FGlc-6P stalls its conversion to Fru-6P, impacting both ATP generation via glycolysis (with potential consequences for cell proliferation) and UDP-GlcNAc synthesis, leading to impaired glycan branching and secretion of the Man5 mAb Fc glycoform.

We further propose that, due to its similarity to Fru-6P, 2FGlc-6P may competitively inhibit PGI (EC 5.3.1.9) and thereby stall glycolysis. This would be consistent with the compromised proliferation observed when cells were fed 2FG at the start of culture (Supplementary Figure 1), given that CHO cells exhibit high glycolytic flux during exponential growth (Ahn and Antoniewicz, 2011), and offers a plausible explanation for why the full 2FG dose on day 0 impairs growth, while delaying the bulk of the dose to later timepoints has a milder impact.

Together, these considerations indicate that 2FG feeding should not begin before day three of culture to protect proliferation and titre. Achieving robust galactosylation control across a broad dose range therefore requires balancing sufficient intracellular UDP-2FGal accumulation (for galactosylation control) against the accumulation of UDP-2FGlc (site occupancy) and 2FGlc-6P (Man5 production and growth impairment).

A key difference between the DP12 C-/GT+ and VRC01 C-/GT+ cell lines was their VCD profiles. DP12 reached a peak cell density of 2.6×10^6^ cells/mL, while VRC01 reached 9.0×10^6^ cells/mL (Figure 1A and Figure 2A). We reasoned that, by being distributed across considerably more cells, considerably lower 2FG accumulation occurred within the VRC01 cells, and that this caused their narrower galactosylation control range.

To address this, we identified that a path forward was to perform 2FG feeding as a cell-normalised dose and not a volume-normalised dose. In parallel, we reasoned that lower peak intracellular 2FG accumulation might reduce production of undesired aglycosylated and Man5 glycans. Considering both aspects, we hypothesised that a higher-frequency, low-dose 2FG feeding strategy (defined on a per-cell basis) would achieve a broader control range while reducing production of undesired aglycosylated and Man5 glycoforms.

To validate the designed feeding strategy, VRC01 C-/GT+ cells were fed 2FG on a per-cell basis across days 3, 4, and 5 of culture, maintaining a constant 2FG dose (normalised to VCD) ranging from 0 to 80 pg/cell. Figure 4 presents the results obtained with the enhanced 2FG feed, where a more pronounced effect on VCD was observed relative to the earlier feeding regimes, with VCD decreasing as 2FG dose increased. However, the maximum VCD reduction was only 16% ± 1% relative to the unfed control (Figure 4A). Although modest, the reduction in VCD is likely attributable to the same glycolysis inhibition effect described previously. 2FG had no discernible impact on q_p_ (table inlay in Figure 4B), and the reduced VCD had only a minor effect on mAb titre – a maximum reduction of 9% ± 1% (Figure 4B).

**Figure 4.**
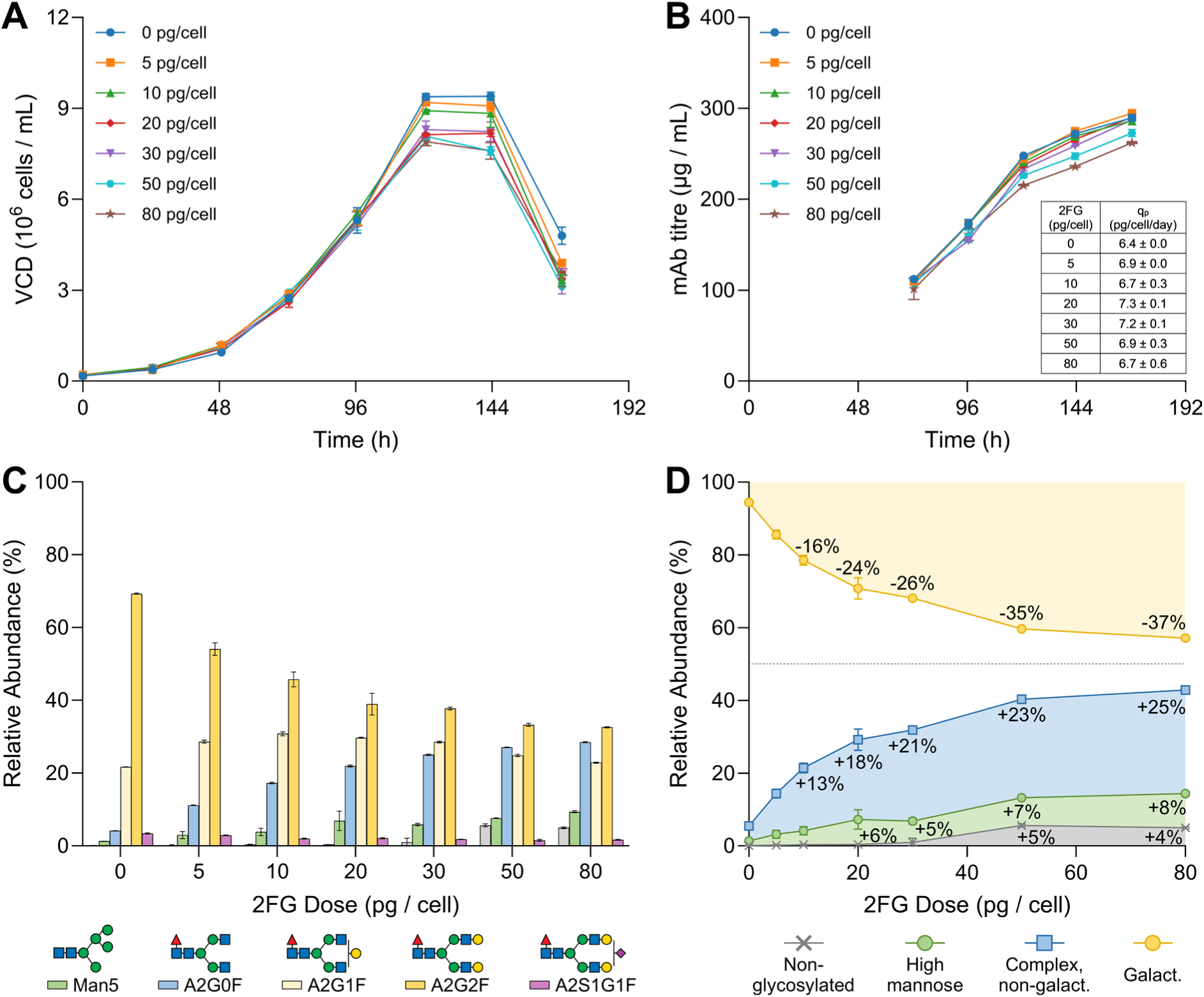
Culture performance of enhanced 2FG feeding on VRC01 C-/GT+ cells. Viable cell density (A), mAb titre (B), Fc glycan profile (C), and Fc glycan motifs (D) are shown for CHO VRC01 C-/GT+ cells treated with the enhanced 2FG feed (0–80 pg/cell), administered across days 3, 4, and 5 of culture. Values represent means ± standard deviation of biological replicates (N = 2). Glycans in (C) follow CFG notation (Varki et al., 2015). In (D), yellow denotes the sum of galactosylated glycans, blue the complex, non-galactosylated A2G0F glycan, green Man5, and grey aglycosylated mAb Fc. Negative values denote losses in galactosylation while positive ones indicate the corresponding increase in non-galactosylation.

Enhanced 2FG feeding produced a dose-dependent reduction in Fc β4-galactosylation, where A2G2F decreased from 69.3% ± 0.2% (unfed) to 32.6% ± 0.1% (80 pg/cell 2FG), while the non-galactosylated species A2G0F increased from 4.2% ± 0.1% (unfed) to 28.5% ± 0.2% (80 pg/cell 2FG) (Figure 4C). Total Fc β4-galactosylation decreased from 94.4% ± 0.1% (unfed) to 57.1% ± 0.2% (80 pg/cell 2FG) (Figure 4D). Of this reduction, 24.8% ± 0.1% was attributable to the increase in A2G0F, 8.2% ± 0.2% to Man5, and 4.3% ± 0.2% to aglycosylated Fc glycoforms (Figure 4D).

Compared with the 10:90 split feed in VRC01 C-/GT+ cells, the enhanced per-cell feeding regime substantially broadened the achievable galactosylation control range, while also maintaining Man5 and aglycosylated glycoform levels considerably lower than those observed in DP12 C-/GT+ cells. These results confirm that adapting the feeding strategy on a per-cell basis and including a third bolus feed is an effective approach to achieving broader, more consistent galactosylation control across cell lines. Nonetheless, some production of Man5 and aglycosylated species persisted at the higher end of the 2FG dose range, indicating that further refinement of the feeding strategy could still simultaneously improve control range and reduction of deleterious mAb Fc glycoforms.

### 3.5. Comparing all three 2FG feeding regimes

To make a quantitative comparison between the 10:90 split feed in DP12 C-/GT+ and VRC01 C-/GT+ cells with the enhanced feeding regime in VRC01 C-/GT+ cells, the average exposure to 2FG, on a per-cell basis, was computed for all three feeding regimes. Using the equations presented in Supplementary Section 3, the average cell exposure to 2FG was obtained by dividing the total amount of 2FG fed throughout culture by the integral average of viable cells from when the first 2FG feed was performed until the end of culture.

Supplementary Figure 2A shows that the split feed in DP12 C-/GT+ and the enhanced feed in VRC01 show comparable average 2FG exposures, with DP12 C-/GT+ ranging from 8 to 168 pg/cell and VRC01 C-/GT+ spanning 13 to 207 pg/cell. The discrepancy between the intended per-cell dose range (5-80 pg/cell) and the computed average exposure arises for two reasons. First, average exposure is calculated only from the time of the first 2FG feed (72 h) onward, whereas the target dose is defined independently of feed timing. Second, the target dose for each feed is calculated based on the VCD present in culture at that specific feeding time point, without accounting for the cumulative effect of prior feeds, whereas the average exposure metric integrates 2FG addition and cell number across the entire post-feeding period.

At lower 2FG doses (0.1 to 0.5 mM), the 10:90 split feed in DP12 C-/GT+ has lower average 2FG exposure than the enhanced feed in VRC01 C-/GT+ cells. For the higher 2FG doses (30 to 80 pg/cell), the enhanced feed in VRC01 C-/GT+ results in a higher average exposure than in DP12 C-/GT+ 10:90 split feed cultures (Supplementary Figure 2A).

Average 2FG exposure for the VRC01 C-/GT+ split feed cultures is considerably lower than in the DP12 split feed and VRC01 enhanced feed. This comparably lower 2FG exposure underlies the narrower galactosylation control range and negligible impacts on VCD and titre obtained in these cultures (Figure 2). Interestingly, the average 2FG exposure for the 2.0 mM split feed and the 20 pg/cell enhanced feed in VRC01 C-/GT+ cells is similar (54 pg/cell and 52 pg/cell, respectively). Supplementary Figure 3 compares culture performance of the two feeding regimes and shows that the enhanced feed has a slightly lower peak VCD, a marginally lower mAb titre, and no significant differences in mAb Fc glycans.

The 2FG feeding profiles compared in Supplementary Figure 3D show close similarity until 96 h of culture, which coincides with the period during which VCD is nearly identical across both cultures. At 96 hours, the enhanced feeding strategy adds a second bolus feed which likely causes the decrease in peak VCD. At 120 hours (day 5), the split feed regime adds a large 2FG bolus, while the enhanced regime adds a much lower one. This comparison shows that the different feeding regimes can achieve similar galactosylation tuning and that feed timing and cumulative 2FG addition may, dose-dependently, impact growth and product titre. Beyond making 2FG dosing transferrable across cell lines, the enhanced feeding regime has three further advantages: (i) it can be adapted based on VCD as cell culture progresses, (ii) it requires less total 2FG to be added, and (iii) it avoids high 2FG doses that may underlie the production of deleterious aglycosylation and Man5 glycans observed in split-fed DP12 C-/GT+ cells.

### 3.6. Ambr^®^ 250 bioreactor runs

Having established the advantages of the split-feed regime, we sought to assess the performance of 2FG feeding in a controlled bioreactor environment and under fed-batch operation. Due to cost, only two FG doses, 5 pg/cell and 15 pg/cell were evaluated. Due to procurement lead time issues, the culture media used in the Ambr^®^ 250 bioreactor runs was changed from ActiPro^TM^, which was used in the shake flask studies, to BalanCD CHO Growth A. Due to this, comparisons between shake flasks and Ambr^®^ 250 runs are made with caution.

All three cultures present nearly identical growth up to 120 hours of culture, after which growth decreases with increasing 2FG dose (Figure 5A). mAb titre is also very similar until 120 hours of culture, after which reductions were observed with increasing 2FG dose (Figure 5B). When compared with unfed cells, decreased mAb titre arose through the combined effect of lower VCD and lower q_p_ (table inlay in Figure 5B). Due to the culture media swap, the Ambr^®^ 250 bioreactor runs achieved a considerably lower peak VCD, when compared to the shake flask studies (Figure 4A). Notably, however, no discernible differences are observed in q_p_, between shake flasks and bioreactors except for the unfed control, which in bioreactors achieved a 42.2% ± 13.3% higher q_p_ than in shake flasks. This variation in q_p_ may also be due to the change in culture media and switch to fed batch operation.

**Figure 5.**
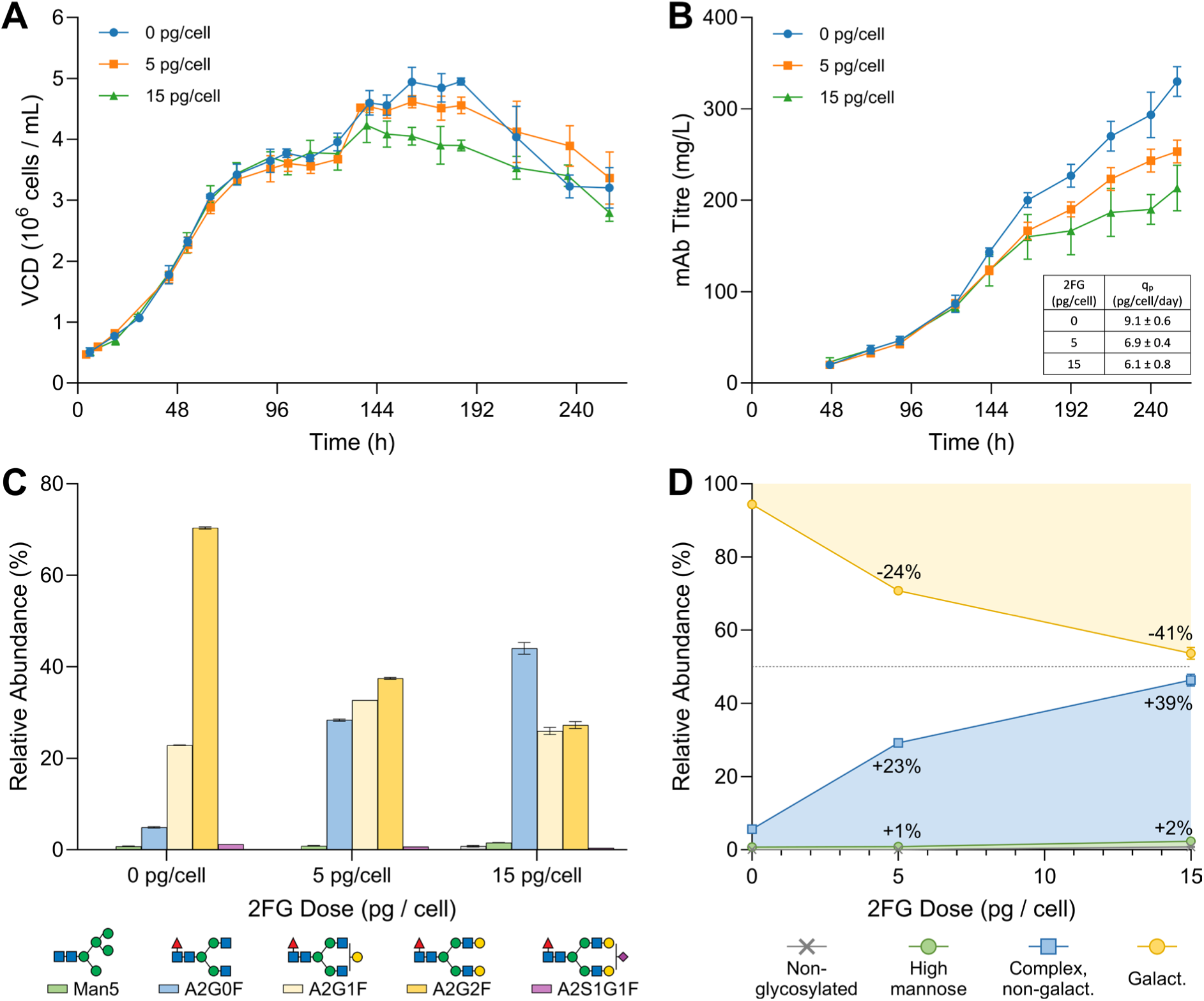
Performance of enhanced 2FG feeding with VRC01 C-/GT+ cells cultured in Ambr^®^ 250 bioreactors. Viable cell density (A), mAb titre (B), Fc glycan profile (C), and Fc glycan motifs (D) are shown for CHO VRC01 C-/GT+ cells cultured in Ambr® 250 bioreactors and treated with the enhanced 2FG feed (0, 5, and 15 pg/cell), administered across days 3, 4, 5, 7, and 9 of culture. Values represent means ± standard deviation of bioreactor replicates (N = 2). Glycan species in (C) are labelled according to CFG notation (Varki et al., 2015). In (D), yellow indicates the sum of galactosylated glycans, blue the A2G0F complex glycan, green the Man5 glycan, and grey aglycosylated mAb Fc. Negative values represent losses in galactosylation, whereas positive values represent the corresponding increase in non-galactosylated species.

For the Ambr^®^ 250 bioreactor runs, glucose, lactate, glutamine, glutamate, ammonia, pH, pCO_2_, pO_2_, Na^+^, K^+^, Ca^2+^, and osmolality were monitored with a Nova BioProfile^®^ FLEX2 cell culture analyser. Time profiles for all culture parameters are presented in Supplementary Figure 4, where only minor differences are observed across 2FG doses in pO_2_, glutamine, glutamate, and osmolality after approximately 168 hours of culture. Glutamine and glutamate presented increasing accumulation with higher 2FG dose. pO_2_ and osmolality also increased with higher 2FG doses. These effects correlate with the lower VCD observed with increasing 2FG doses (Figure 5A).

Figure 5C shows the mAb Fc glycoprofiles obtained in the Ambr^®^ 250 bioreactor runs. Compared with the shake flask cultures (Figure 4C), these runs achieved a much more pronounced reduction in galactosylation at considerably lower 2FG doses and without production of deleterious aglycosylated and Man5 glycoforms. The 15 pg/cell 2FG dose yielded a 41% reduction in galactosylation with only a 2% increase in Man5 production (Figure 5D), reflecting a direct shift from A2G1F and A2G2F to A2G0F, the ideal scenario for galactosylation control.

Supplementary Figure 5 compares the major mAb Fc glycosylation motifs (galactosylation, aglycosylation, high mannose, and complex non-galactosylated) across the DP12 C-/GT+ split feed, the VRC01 C-/GT+ enhanced feed, and the VRC01 C-/GT+ Ambr^®^ 250 runs. The fraction of glycan motifs obtained with the highest 2FG exposure of each feeding regime are summarised in Table 1 and show that the DP12 split feed achieves the highest galactosylation reduction (2.1-fold) at an average 2FG exposure of 168.3 pg/cell but at the expense of large increases in aglycosylation and high mannose glycans. The Ambr^®^ 250 run, with a nearly 10-fold lower 2FG exposure, achieves a 1.8-fold reduction in galactosylation and low levels of aglycosylation and high mannose glycans, an outcome that could not be predicted from the shake flask data alone. Understanding the mechanisms that drive the higher 2FG sensitivity in the Ambr^®^ 250 runs is essential to rationally guide further optimisation of the feeding strategy.

**Table 1.** Comparison of glycan motifs at the highest 2FG dose for each feeding regime.

|  | **DP12**<br>**Split** | **VRC01**<br>**Split** | **VRC01**<br>**Enhanced** | **VRC01**<br>**Ambr<sup>®</sup> 250** |
| --- | --- | --- | --- | --- |
| Max. 2FG exposure (pg/cell) | 168.3 | 54.3 | 207.1 | 18.56 |
| Galactosylation (%) | 45.9 ± 0.2 | 71.8 ± 1.1 | 57.1 ± 0.2 | 53.7 ± 1.6 |
| Galactosylation fold-change | -2.1 ± 0.0 | -1.3 ± 0.0 | -1.5 ± 0.0 | -1.8 ± 0.1 |
| Aglycosylation (%) | 12.6 ± 0.7 | 0.3 ± 0.1 | 5.0 ± 0.2 | 0.8 ± 0.2 |
| High mannose (%) | 26.8 ± 0.4 | 5.7 ± 0.2 | 9.4 ± 0.3 | 1.5 ± 0.1 |
| Complex Non-galact. (%) | 14.6 ± 0.5 | 21.7 ± 0.7 | 28.5 ± 0.2 | 44.0 ± 1.3 |

### 3.7. Glycoforms produced post 2FG feeding

To gain a better understanding of how the different feeding regimes shape the obtained glycosylation profiles, the fraction of mAb glycoforms secreted after 2FG feeding were calculated with Equation 1. Conceptually, the post-feed secreted glycoform distributions represent the glycosylation profile that the cells would produce after the first 2FG feed to go from the profile generated by the unfed controls to the measured endpoint glycan distribution.

Split feed DP12 C-/GT+ cells (Figure 6A) showed considerable aglycosylation and Man5 production, even at relatively low average 2FG exposure. Split feed VRC01 C-/GT+ cells (Figure 6B) showed a considerable dose-dependent reduction in A2G2F, with concomitant accumulation of A2G1F and A2G0F, some Man5 production, but undetectable aglycosylation. Enhanced feed VRC01 C-/GT+ cells (Figure 6C) produced a marked reduction in A2G2F with a concomitant increase in A2G1F at lower average 2FG exposure. With increasing 2FG levels, A2G2F and A2G1F both decreased to yield A2G0F, with low but significant levels of aglycosylated and Man5 glycoforms. In the Ambr^®^ 250 runs with VRC01 C-/GT+ cells (Figure 6D), only marginal amounts of aglycosylation and Man5 were produced, with a considerable shift from the galactosylated A2G1F and A2G2F species to A2G0F occurring even at considerably lower 2FG exposure than in the other cultures.

**Figure 6.**
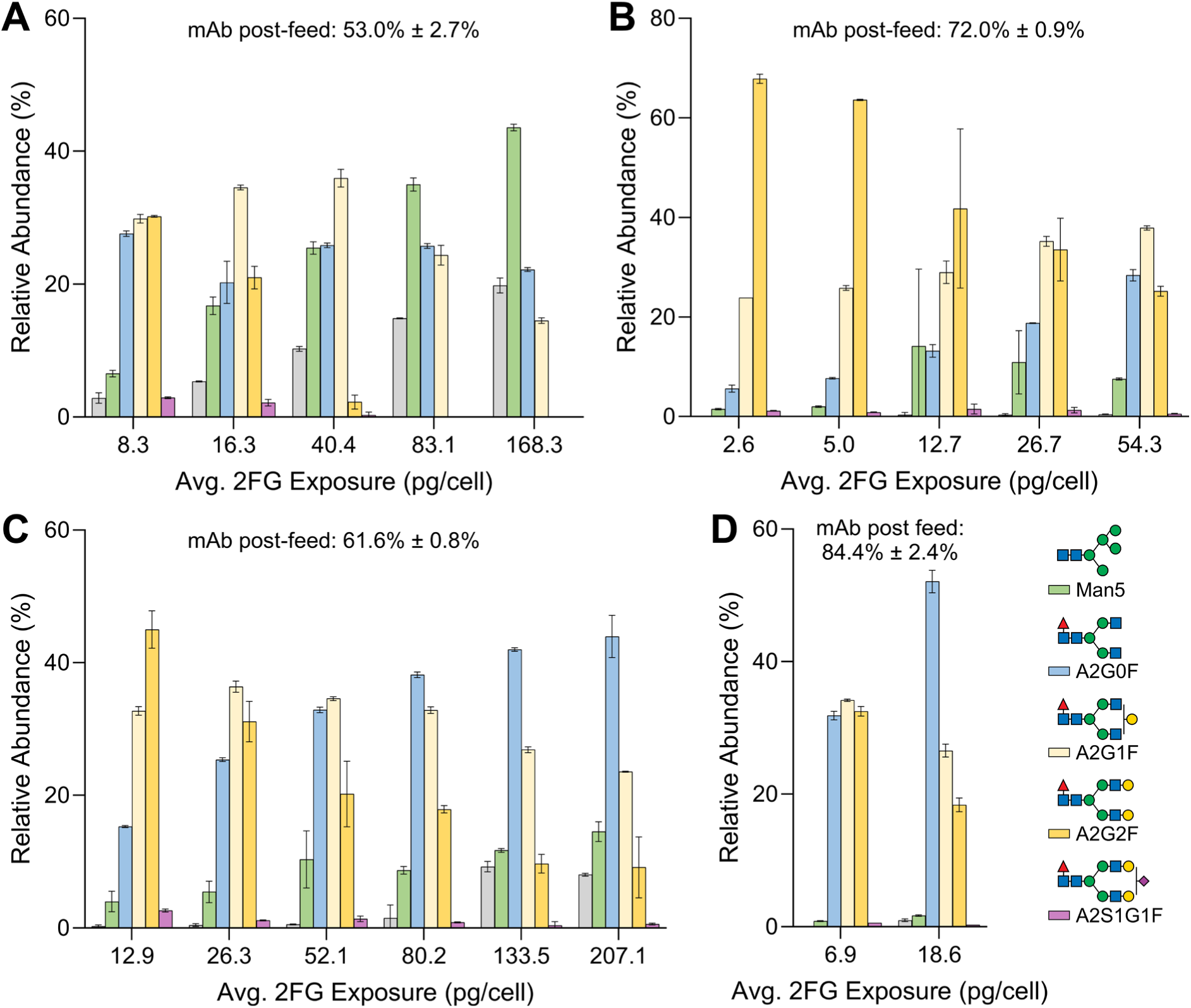
Glycoform distributions secreted after 2FG feeding. Post-feed glycoform distributions for the DP12 C-/GT+ split feed (A), VRC01 C-/GT+ split feed (B), VRC01 C-/GT+ enhanced feed (C), and Ambr^®^ 250 bioreactor (D) cultures are shown. These were computed with Equation 1 using the measured endpoint glycoprofiles and their respective unfed controls as inputs. Calculations were performed for each biological replicate, and the values represent means ± S.D. across the replicates (N=2). Labels above each panel indicate the proportion of mAb produced after the first 2FG feed. Glycan species are labelled according to CFG notation (Varki et al., 2015).

Crucially, the post-feed secreted glycoforms implicitly reflect the impact of mAb production dynamics on the effectiveness of the 2FG feeding strategies: when large amounts of mAb are produced before 2FG feeding, the glycoforms produced after 2FG feeding will be diluted by those that are present under unfed conditions, resulting in a narrower control range that cannot be broadened with higher 2FG doses because this would likely cause production of deleterious aglycosylated and Man5 glycoforms, as seen in DP12 C-/GT+ split feed cultures. Conversely, when a large proportion of mAb is secreted after 2FG feeding, the dilution effect will be reduced and the endpoint glycosylation profile will be more heavily influenced by 2FG exposure, even at lower 2FG doses.

Building on this reasoning, Figure 6 also reports the proportion of mAb produced after the first 2FG feed at the top of each panel, allowing the extent of this dilution effect to be directly assessed for each feeding regime. For DP12 C-/GT+ split feed cultures, only 53% of the total mAb was produced after 2FG feeding. As a result, even though the lowest 2FG dose (8.3 pg/cell) produced a substantial post-feed reduction in galactosylated species, from 95% in the unfed control to approximately 60% post-feed, this translated into only a modest reduction, to around 80%, in the endpoint glycoprofile. In contrast, the Ambr^®^ 250 runs, conducted in fed-batch mode over 264 hours, achieved a much higher post-feed production fraction, with 84% of total mAb produced after the first 2FG feed. As a result, the post-feed reduction in galactosylated species was closely reflected in the endpoint glycoprofile: at 18.6 pg/cell 2FG exposure, cells produced 45.2% ± 2.1% galactosylated Fc glycoforms post-feeding, translating into an endpoint reduction from 94.3% ± 0.2% (unfed control) to 53.7% ± 1.6%, with minimal dilution from pre-feed mAb glycoforms. Together, these two cases illustrate how the proportion of mAb secreted after 2FG feeding directly determines the extent to which post-feed glycosylation changes are reflected in the final, measurable glycoprofile.

Taken together, these results indicate that the achievable galactosylation control range correlates with the proportion of mAb secreted after 2FG feeding: in cultures where most mAb is produced post-feed, as in the Ambr® 250 runs, low 2FG doses that avoid triggering aglycosylation or Man5 production are sufficient to regulate galactosylation across a broad range, whereas in cultures where most mAb is produced before feeding, as in DP12, achieving a comparable control range would require substantially higher 2FG doses, with a concomitant risk of producing these deleterious glycoforms. Although the obtained results are consistent with this framework, the change in culture media may also have an effect.

### 3.8. Impact of glycosylation profiles on CD16a (FcγIIIa) binding

Antibody-dependent cell-mediated cytotoxicity (ADCC) is a key effector function of therapeutic mAbs, and its efficacy is modulated, in large part, by the affinity of the Fc region for CD16a (FcγRIIIA) expressed on natural killer cells (Jefferis, 2005; Nimmerjahn and Ravetch, 2008). Fc N-glycosylation, and in particular the extent of galactosylation and the presence of core fucosylation, has been widely reported to influence this interaction, with hypergalactosylated glycoforms associated with enhanced CD16a binding (Subedi and Barb, 2016; Thomann et al., 2016) and presence of core fucose reducing CD16a affinity (Shinkawa et al., 2003). Aglycosylated Fc is associated with abolished binding (Krapp et al., 2003; Tao and Morrison, 1989), and high mannose mAb glycoforms, including Man5, have been shown to enhance ADCC (Zhou et al., 2008) due to absence of fucosylation, although they are still outperformed by complex afucosylated glycoforms (Kanda et al., 2007), which do not carry risks of reduced serum half-life (Baumeister et al., 2026) and immunogenicity (Dasgupta et al., 2007; Delignat et al., 2020).

Because the 2FG feeding strategy described here enabled production of a broad and well-characterised range of mAb Fc glycoforms, spanning highly galactosylated species through to elevated levels of non-galactosylated, high-mannose, and aglycosylated forms, this platform offered a valuable opportunity to directly examine the relationship between Fc glycosylation profile and CD16a binding affinity. To this end, Bio-Layer Interferometry (BLI) assays were performed on selected samples spanning this glycosylation range to quantitatively assess CD16a binding.

BLI results for the 0, 0.2, and 2.0 mM 2FG doses in DP12 C-/GT+ split feed cultures, and the 0, 10, and 20 pg/cell doses in the enhanced 2FG feed, are presented in Figure 7A (representative sensorgrams are shown in Supplementary Figure 6). The obtained equilibrium dissociation constants (*K_D_*) ranged between 30 and 60 nM across all samples, consistent with the reported affinity range for FcγRIIIA (*K_D_* ≈ 10-400 nM) (Van Coillie et al., 2022) and closely matching values previously reported for CD16a binding to IgG1 Fc glycoforms bearing complex-type N-glycans (Subedi and Barb, 2018). Across both mAbs, *K_D_* increased with increasing 2FG dose, indicating a progressive reduction in CD16a binding affinity with changes in mAb Fc glycosylation.

**Figure 7.**
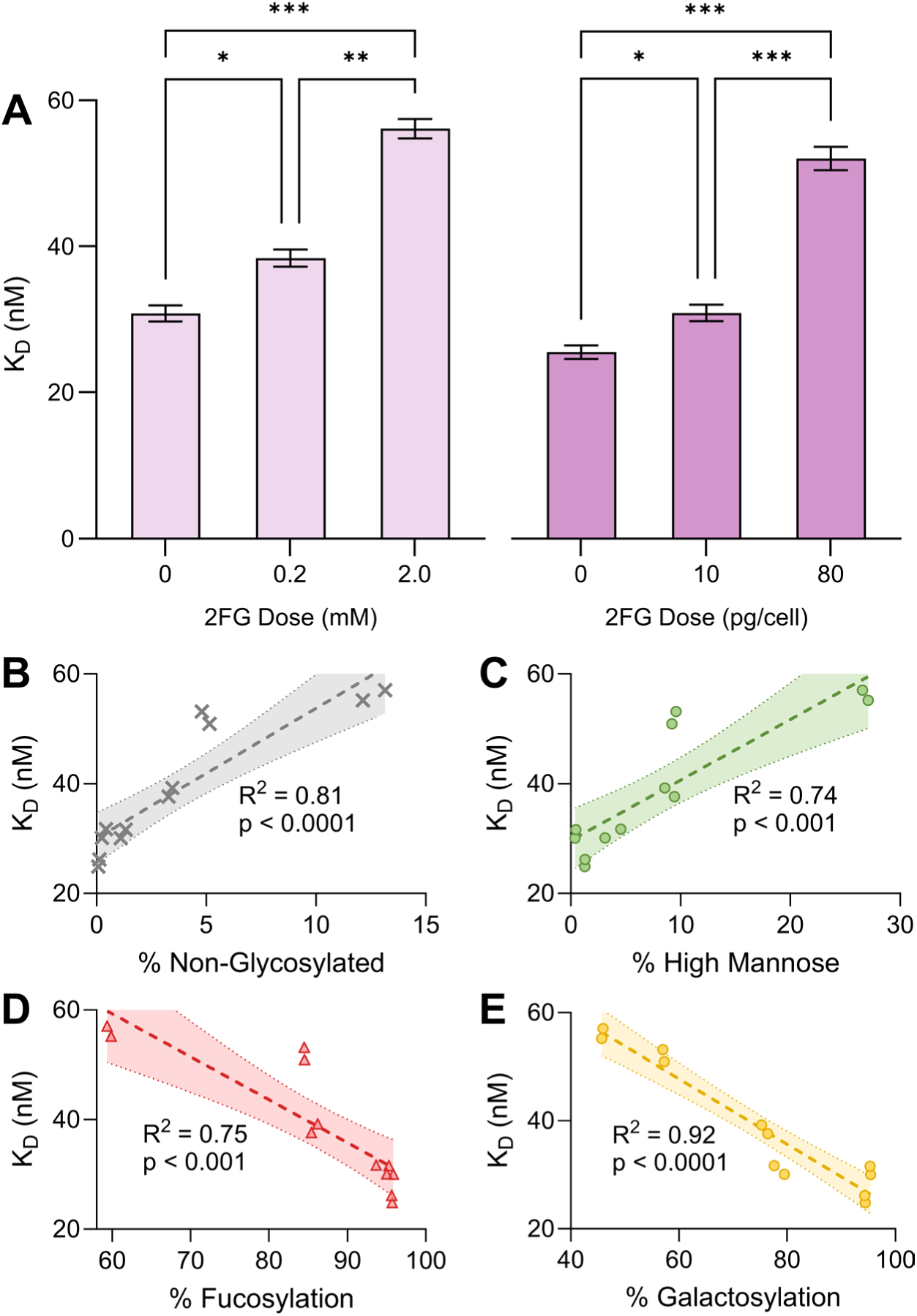
CD16a (FcγRIIIA) binding affinity across 2FG feeding regimes. (A) Equilibrium dissociation constants (*K_D_*) obtained by BLI for the tested 2FG doses across feeding regimes, shown for DP12 C-/GT+ split feed (light purple) and VRC01 C-/GT+ enhanced feed (dark purple). Bars represent mean values ± standard deviation of technical replicates. One-way ANOVA with Tukey’s post hoc test (GraphPad Prism 11.0.2) was performed to assess differences between values (*p < 0.05, **p < 0.01, ***p < 0.001, ****p < 0.0001). (B–E) Correlation and linear regression analyses between *K_D_* and the fraction of each glycan motif across all samples and feeding regimes: aglycosylation (B), high mannose (C), fucosylation (D), and galactosylation (E). Shaded areas represent the 95% confidence intervals of the fitted regression lines. The coefficient of determination (*R*^2^) and the significance of the slope are inlaid in each panel. Linear regression analysis was performed using GraphPad Prism 11.0.2.

Because the samples exhibit variable extents of several glycan motifs (aglycosylation, high mannose, fucosylation, and galactosylation), each of which has been reported to influence CD16a binding affinity, we sought to identify which of these motifs were the key contributors to the observed reductions in binding affinity. To this end, we performed correlation analysis, comparing the fraction of each glycan motif across all six antibody samples against their measured *K_D_*s (Figure 7B through Figure 7E). Aglycosylation showed a positive correlation with *K_D_*, consistent with the loss of Fc effector function previously reported for aglycosylated IgG (Krapp et al., 2003; Tao and Morrison, 1989). High mannose glycoforms also correlated positively with *K_D_*. While several studies report that Man5 glycoforms bind CD16a more tightly than fucosylated complex glycans due to the inherent absence of core fucose (Zhou et al., 2008), other work using affinity-based methods has reported weaker CD16a binding for high-mannose glycoforms relative to G0F (Kanda et al., 2007), consistent with our observations.

Intriguingly, fucosylation correlated negatively with *K_D_*, indicating that higher fucosylation was associated with increased, rather than decreased, binding affinity – the opposite of what is widely reported (Shields et al., 2002; Shinkawa et al., 2003). It is possible that intermediate levels of fucosylation (as opposed to complete afucosylation) underlie this discrepancy. Increased galactosylation showed a negative correlation with *K_D_*, which is consistent with previous reports that galactosylation enhances CD16a binding affinity (Subedi and Barb, 2018; Thomann et al., 2016). Supplementary Figure 7 shows the correlation matrix for *K_D_* and all glycan motifs. Notably, all four glycosylation motifs were, themselves, correlated with one another: increases in aglycosylation and high mannose content were closely accompanied by reductions in fucosylation and galactosylation. These close correlations may also underlie the unexpected direction of the fucosylation correlation with *K_D_*.

Although the correlation analysis (Figures 7B through 7E) indicates that all four glycosylation motifs have statistically significant linear correlations (p < 0.001) with *K_D_*, the confidence bands for the fitted lines vary considerably among motifs, as does the coefficient of determination (*R*^2^). High mannose and fucosylation show the broadest confidence intervals and lowest *R*^2^ values, whereas galactosylation shows the tightest confidence intervals and the highest *R*^2^ (0.92), suggesting that galactosylation is the primary driver of the enhanced binding affinity observed across these glycan distributions.

## 4. Conclusions

This work demonstrates that 2-deoxy-2-fluorogalactose (2FG) feeding provides a robust and tuneable strategy for controlling mAb Fc galactosylation when combined with the C-/GT+ hypergalactosylating cell engineering platform to yield a broad spectrum of glycosylation states. In DP12 C-/GT+ cells, the 10:90 split-feed regime achieved a galactosylation reduction from 95% to 46%, but this control range came at the cost of substantial aglycosylation (13%) and Man5 production (27%), both of which are undesirable Fc glycoforms with known impacts on effector function, serum half-life, and immunogenicity risk. The same feeding regime produced a markedly narrower control range in VRC01 C-/GT+ cells, revealing that a fixed, volume-normalised 2FG dose is not directly transferable across cell lines with different growth and production dynamics.

To address this, we developed a cell-normalised, multi-bolus feeding strategy that substantially broadened the achievable galactosylation control range in VRC01 C-/GT+ cells while keeping aglycosylation and Man5 production considerably lower than in the original DP12 split-feed regime. Quantitative comparison of average per-cell 2FG exposure across all tested regimes confirmed that cellular exposure, rather than the nominal 2FG dose, is likely the key variable governing the extent and character of shift in glycosylation. We have used this cellular exposure metric to design comparable feeding strategies across cell lines with differing growth kinetics while simultaneously reducing production of aglycosylated and high mannose glycoforms. Analysis of post-feed secreted glycoforms further showed that the proportion of mAb produced after 2FG addition is a critical determinant of the achievable control range: cultures in which most product is secreted after feeding require substantially lower 2FG exposure to achieve a given galactosylation shift and are correspondingly less prone to producing deleterious glycoforms.

This principle was validated in Ambr^®^ 250 bioreactor runs, where fed-batch cultivation extended the post-feed production window and yielded a comparable reduction in galactosylation with, roughly, an order of magnitude lower 2FG exposure than in shake flask cultures. The Ambr^®^ 250 runs also yielded only marginal production of aglycosylated and Man5 glycoforms. Although the different culture media used for Ambr^®^ 250 cultures may also contribute to the broader control range achieved, our results highlight that design of the 2FG feeding strategy should account not only for VCD and 2FG sensitivity, but also for the mAb production kinetics of the culture format in which it will be deployed.

Finally, BLI analysis confirmed that the glycoform diversity generated by 2FG feeding translates into a functionally meaningful range of CD16a (FcγRIIIA) binding affinities, with galactosylation emerging as the strongest correlate of *K_D_* across all tested samples, while aglycosylation, high mannose, and fucosylation showed weaker and, in the case of fucosylation, counterintuitive associations that are likely confounded by the strong inter-correlation among these glycan motifs. Together, these findings establish 2FG feeding, guided by per-cell dosing and production-phase considerations, as a practical and generalisable tool for tuning mAb Fc galactosylation and, by extension, CD16a-mediated effector function, offering a flexible and robust method to explore structure-function relationships during antibody product design.

Future work will focus on validating the robustness of the cell-normalised 2FG feeding strategy across a broader panel of cell lines, media, and bioreactor formats, building on the transferability already demonstrated between DP12 and VRC01 C-/GT+ cells. Direct testing of the proposed model of 2FG metabolism, through intracellular metabolite analysis, should also be performed to further refine the feeding strategy with the goal of broadening the galactosylation control range while minimising aglycosylation and Man5 glycoform production. Our combined metabolic and cellular glycoengineering workflow could also be extended to other therapeutic glycoprotein formats, including bi- and multi-specific antibodies.

## Acknowledgments

AT and IJV gratefully acknowledge funding from Research Ireland through the Solid-State Pharmaceutical Cluster – RI’s Research Centre for Pharmaceuticals (12/RC/2275_P2 SSPC). MG thanks the Irish Research Council (GOIPG/2015/3863) for their financial support. The authors thank the National Vaccine Center at the National Institutes of Health for the CHO VRC01 cell line. The authors thank Dr. Mark Sheehan and Éadaoin O’Donoghue for their kind assistance with the Ambr^®^ 250 bioreactor runs.

## Declaration of generative AI and AI-assisted technologies in manuscript preparation

During preparation of this work, the authors used Anthropic PBC’s Claude Sonnet 5.0 to improve the text’s clarity and conciseness. After using the tool, the authors reviewed and edited the content as needed and take full responsibility for the content of the published article.

## Author contributions

AT: investigation, methodology, data analysis, and writing – original draft. MG: investigation, methodology, and data analysis. IGA: conceptualisation and methodology. KD: investigation, methodology, and data analysis. BG: resources, supervision, and writing – review and editing. SC: investigation, methodology, and data analysis. JB: resources, supervision, funding acquisition, and writing – review and editing. IJV: conceptualisation, data analysis, funding acquisition, project administration, resources, supervision, and writing – review and editing.

