## Supplementary Information for "Next-gen glycoengineering: combining cellular and metabolic engineering to fine-tune mAb β1,4-galactosylation"

### 1. **Fc N-glycan analysis**

50 µg of protein A purified and neutralized sample were brought to a concentration of 1 mg/mL, and enzymatic digestion was initiated by the addition of 1uL of IdeS enzyme (FABricator, Genovis). The digestion was carried out for 2h at 37°C generating non-reduced Fab domain and scFc subunits. Samples were immediately transferred to the autosampler at 5°C to perform LC-MS analysis.

Separation of the Fc from the Fab region was achieved by reversed phase chromatography. The analysis was performed on 1 µL of sample and in triplicate using a Vanquish Horizon UHPLC (Thermo Fisher Scientific, Sunnyvale, CA, US) equipped with a MAbPac RP 2.1 x 50 mm column (Thermo Fisher Scientific, Cat. No. 088648). LC separation was performed via linear gradient using, as mobile phases, water with 0.1 % formic acid (A) and acetonitrile with 0.1 % formic acid (B). The gradient started at 25% B, and this composition was held for 1 minute before starting a gradient to 45% B in 15 minutes. Following the gradient, the column was washed for 2 minutes at 80% B before restoring the starting conditions and re-equilibration for 7 minutes. Flow rate was kept at 0.3 mL/min and column temperature was held at 80 °C. After chromatographic separation, the analyte was introduced to a Thermo Scientific Exploris 240 Orbitrap mass spectrometer equipped with an OptaMax NG source (Thermo Fisher Scientific, Sunnyvale, CA, US). MS settings were as follows: spray voltage 3.8 kV, sheath gas 25 arbitrary units (AU), auxiliary gas 10 AU, ion transfer capillary temperature 320 °C, and vaporizer temperature 150 °C. Analysis mode was set to Intact Protein using Low Pressure for trap pressure settings. Scan parameters were as follows: resolution setting was 120,000 at 200 m/z, scan range was 600 – 3,000 m/z, RF lens was set to 80%, acquisition gain control was 300% and maximum injection time was 200 ms using 5 microscans. All data were acquired using Chromeleon CDS v 7.3.1 for both instrument control and data acquisition.

Raw data were analysed using BioPharma Finder software v4.1. In the chromatograms, two peaks were present, one corresponding to scFc portion and the other to the Fab but only the scFc peak was analysed further to obtain N-glycan profiling. Deconvolution was performed using the "average over selected time range" option using the Xtract algorithm for isotopically resolved spectra. Output mass range was fixed between 23,500 and 26,500 Da, while charge states range was set between 5 and 50, considering only masses deconvoluted from a minimum of 5 consecutive charge states. Identification was performed using protein sequences for the scFc region, including C-terminus lysine clipping as a fixed modification and N-glycans as side chain variable modifications. Mass accuracy was generally lower than 5 ppm and identification was accepted only for species present in at least 2 replicates. Average MS signal intensity for each identified component was used to calculate fractional abundance.

### 2. Preliminary assessment of 2FG feeding regimes

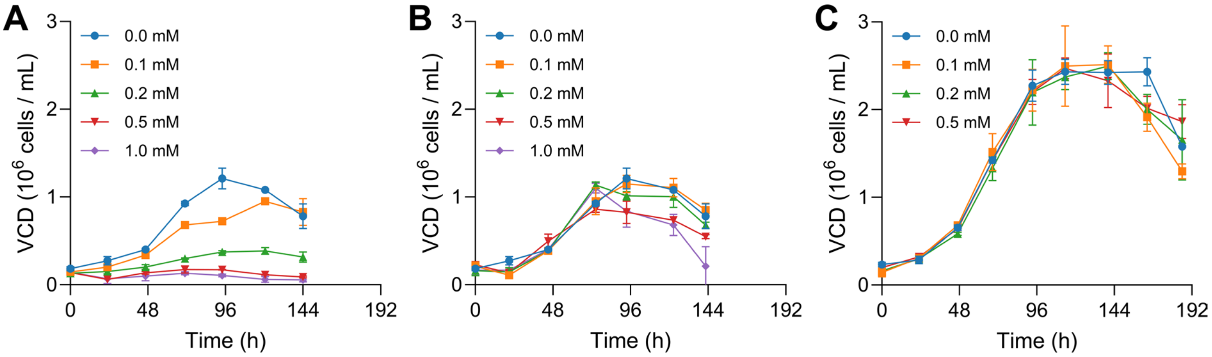


**Supplementary Figure 1.** Cell culture growth curves for the initial test runs where 2dFGal was fed at the beginning of the culture (a), at mid-exponential phase (b) and, 10% on day 3 and 90% on day 5 (c). The cultures presented in (a) and (b) are static in t-25s, while the last run (c) was performed in 125mL shake flasks. Error bars represent the standard deviation between biological replicates (N=2).

### 3. Comparison of 2FG feeding regimes

The average 2FG dose ($\overline{D_{2FG}}$) was computed for the 10:90 split and enhanced feeding regimes using Supplementary Equation S1, where $2FG_{tot,add.}$ is the overall endpoint 2FG concentration and $IAVCD_{FG}$ is post-2FG feed integral-average viable cell density, as computed with Supplementary Equation S2. In Supplementary Equation S2, $IVCD_{FG,0}$ is the integral of viable cell density at the first 2FG feeding timepoint, $IVCD_f$ is the integral of viable cell density at the culture endpoint, $t_{FG,0}$ is the time at which the first 2FG feed was made and $t_f$ is the culture endpoint. Supplementary Equation S3 presents how the IVCD values were computed using the trapezoidal integration rule, where $VCD(t_{k-1})$ and $VCD(t_k)$ are the viable cell densities at culture times $t_k$ and $t_{k-1}$, respectively.

$$\overline{D_{2FG}} = \frac{2FG_{tot,add.}}{IAVCD_{D3}} \qquad \text{Eq. S1}$$

$$IAVCD_{FG} = \left(\frac{IVCD_f - IVCD_{FG,0}}{t_f - t_{FG,0}}\right) \qquad \text{Eq. S2}$$

$$IVCD_k = \left(\frac{1}{2}\right)\sum_{k=1}^{N}[VCD(t_{k-1}) + VCD(t_k)](t_k - t_{k-1}) \qquad \text{Eq. S3}$$

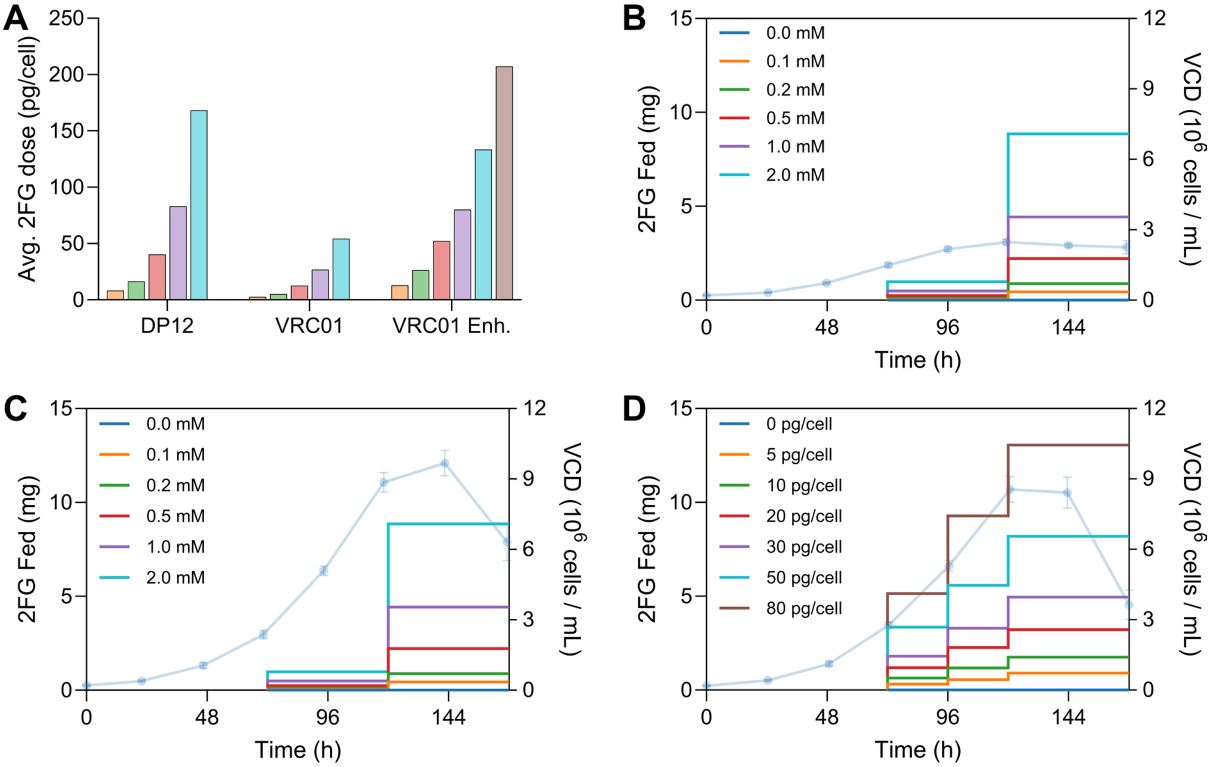


**Supplementary Figure 2. Comparison of 2FG feeding regimes**

(A) Shows the average 2FG dose across the 10:90 split feed for DP12 and VRC01 as well as the enhanced feeding regime (VRC01 Enh.). (B) shows the amount of 2FG fed throughout culture for the 10:90 split feed in DP12 C-/GT+ cells and across 0 to 2.0 mM concentration range. (C) presents the 2FG feed for the 10:90 split feed in VRC01 C-/GT+ cells. (D) shows the enhanced 2FG feeding regime across doses from 0 to 80 pg/cell. (B) through (C) are overlaid on the average VCD profile for each feeding regime (left axis).

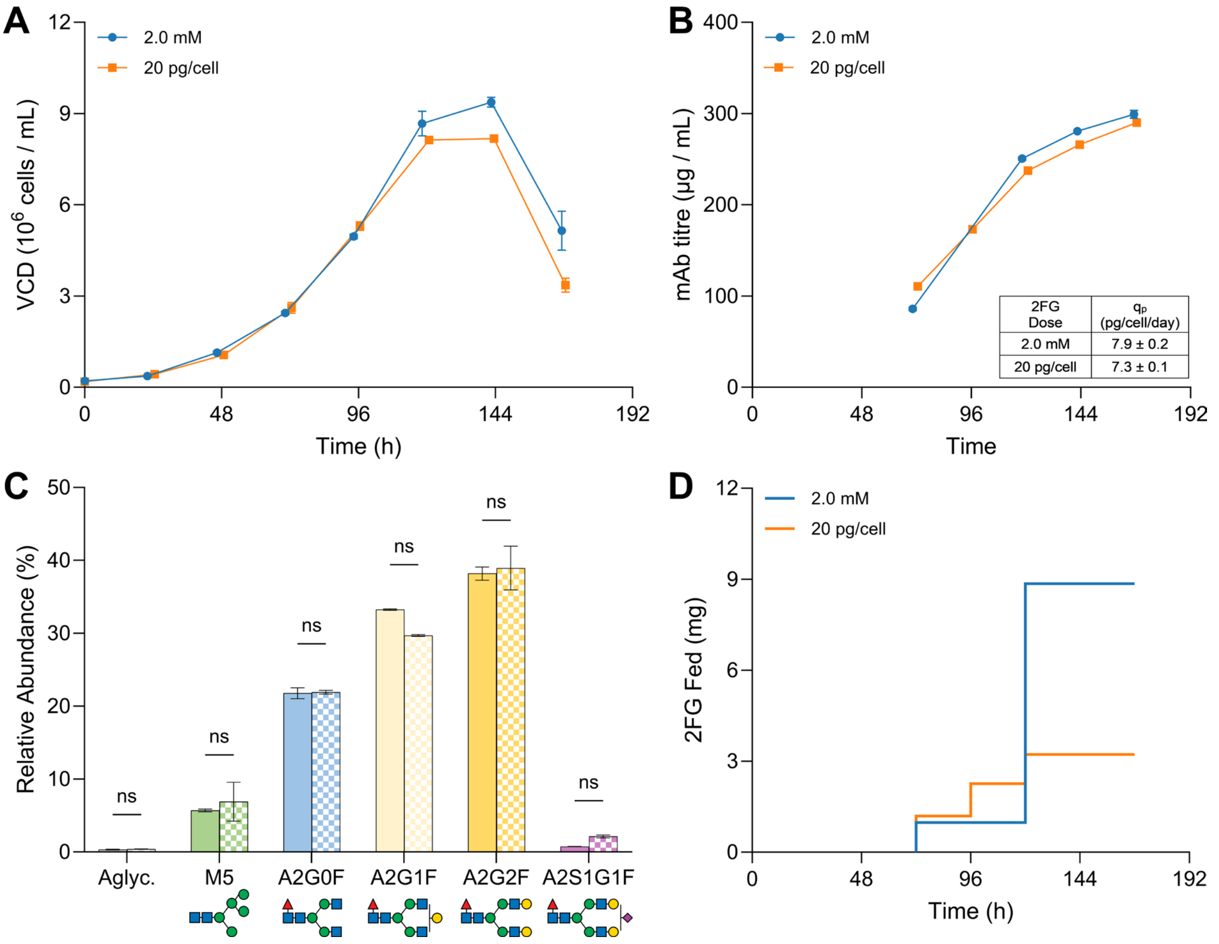


**Supplementary Figure 3. Culture performance comparison**

Comparison of VCD (A), mAb titre (B), glycan profile (C), and 2FG feeding schedule (D) between the 10:90 split feed (blue) and the enhanced feed (orange) regimes. All values correspond to means ± standard deviation of biological replicates (N=2). Glycans in (C) follow CFG notation (Varki et al., 2015). The comparisons shown in (C) correspond to a two-way ANOVA with Tukey's post hoc test (GraphPad Prism 11.0.2). ns indicates significant differences were not observed ($p \geq 0.05$).

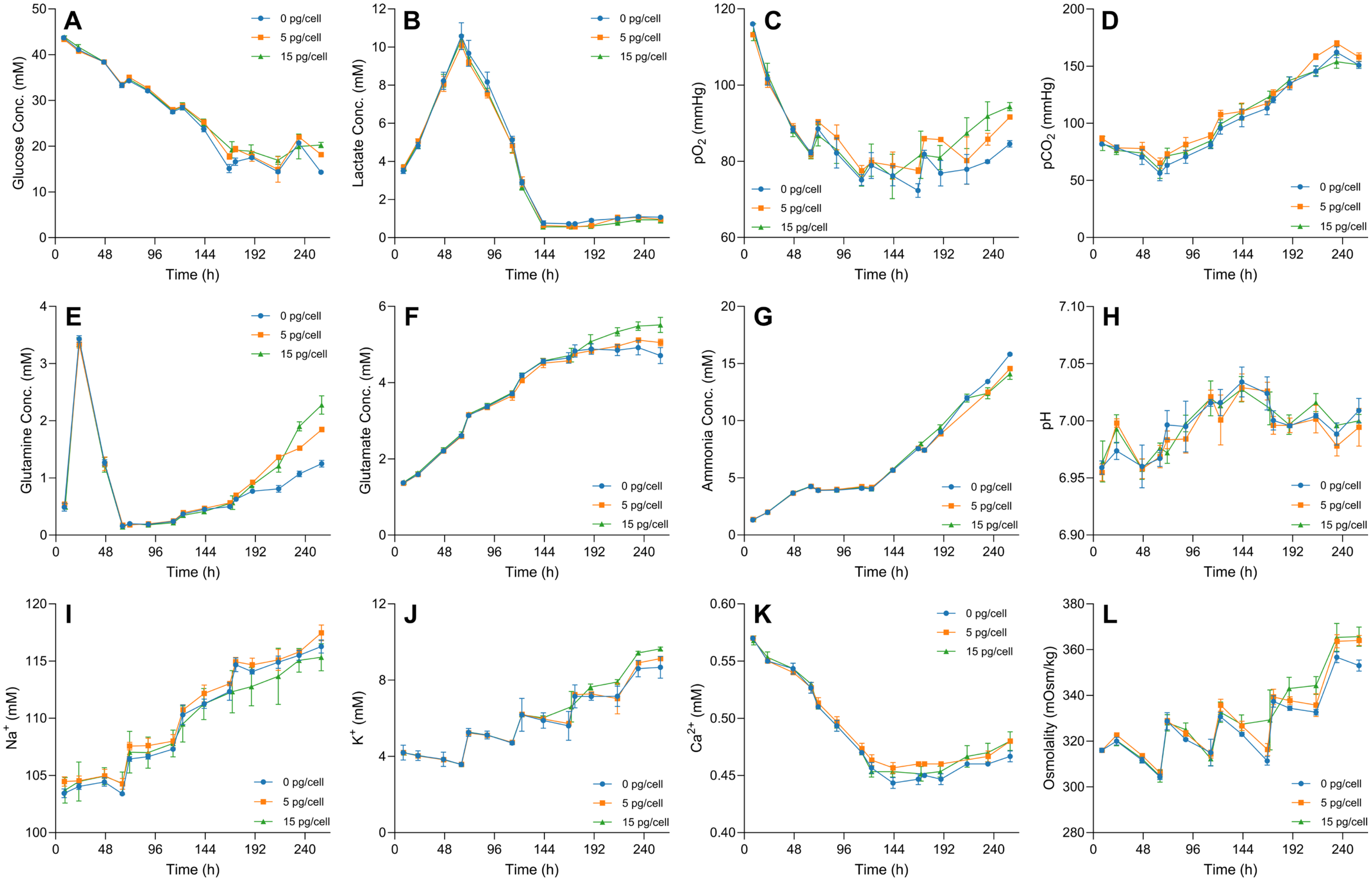


**Supplementary Figure 4. BioProfile® FLEX2 measurements for the Ambr®250 bioreactor runs**

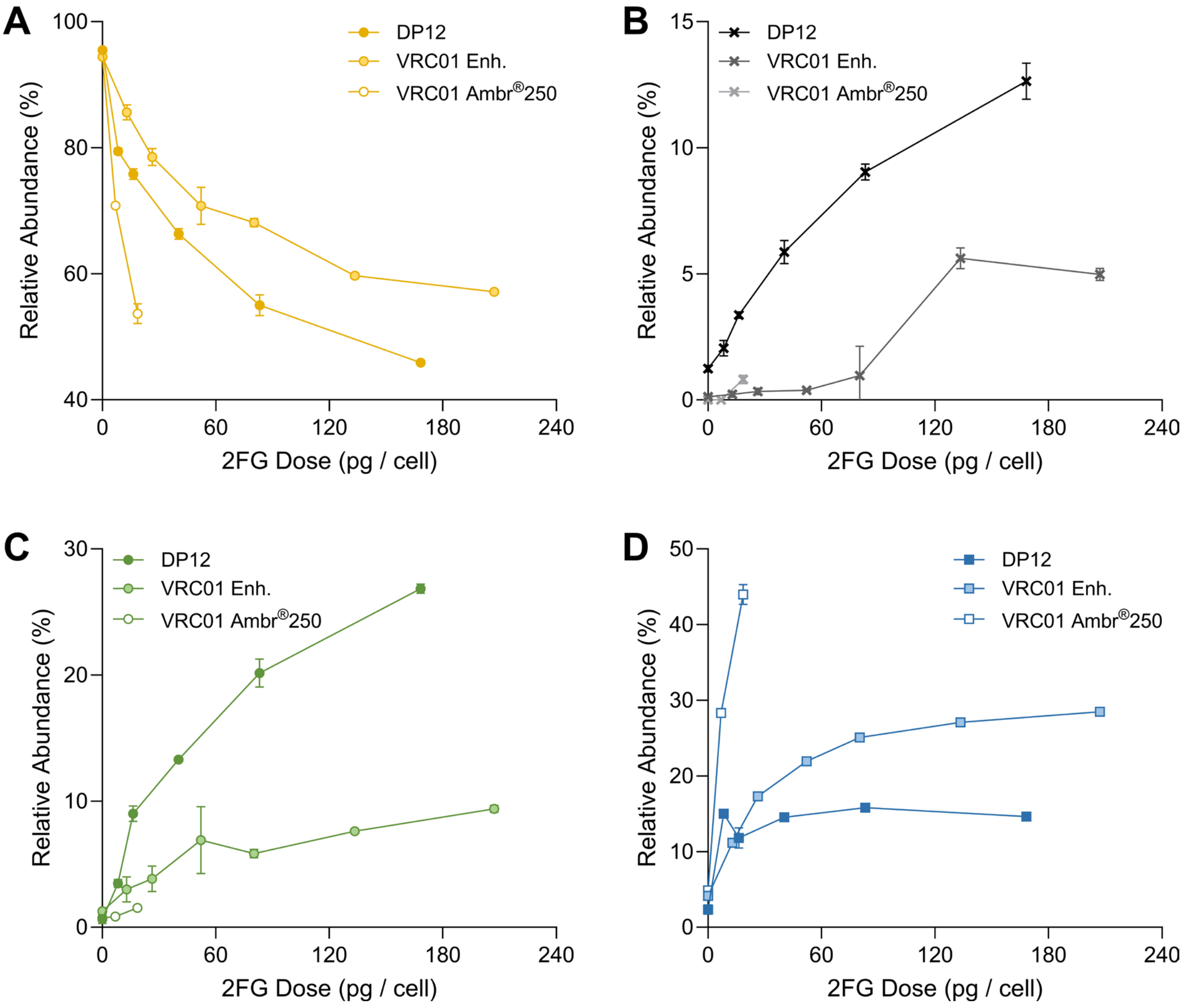


**Supplementary Figure 5. Comparison of main mAb glycosylation motifs produced with DP12 and VRC01 cells cultured under split and enhanced 2FG feeding in shake flasks and Ambr® 250 bioreactors**

Galactosylated (A), aglycosylated (B), high mannose (C), and complex non-galactosylated (D) motifs produced under different 2FG doses are shown. Profiles for DP12 cultured with the 10:90 split fed (DP12), shake flask VRC01 cultures with enhanced feeding (VRC01 Enh.), and VRC01 bioreactor runs with enhanced feeding (VRC01 Ambr®250) are shown. Data corresponds to mean ± S.D. for duplicate cultures (N=2).

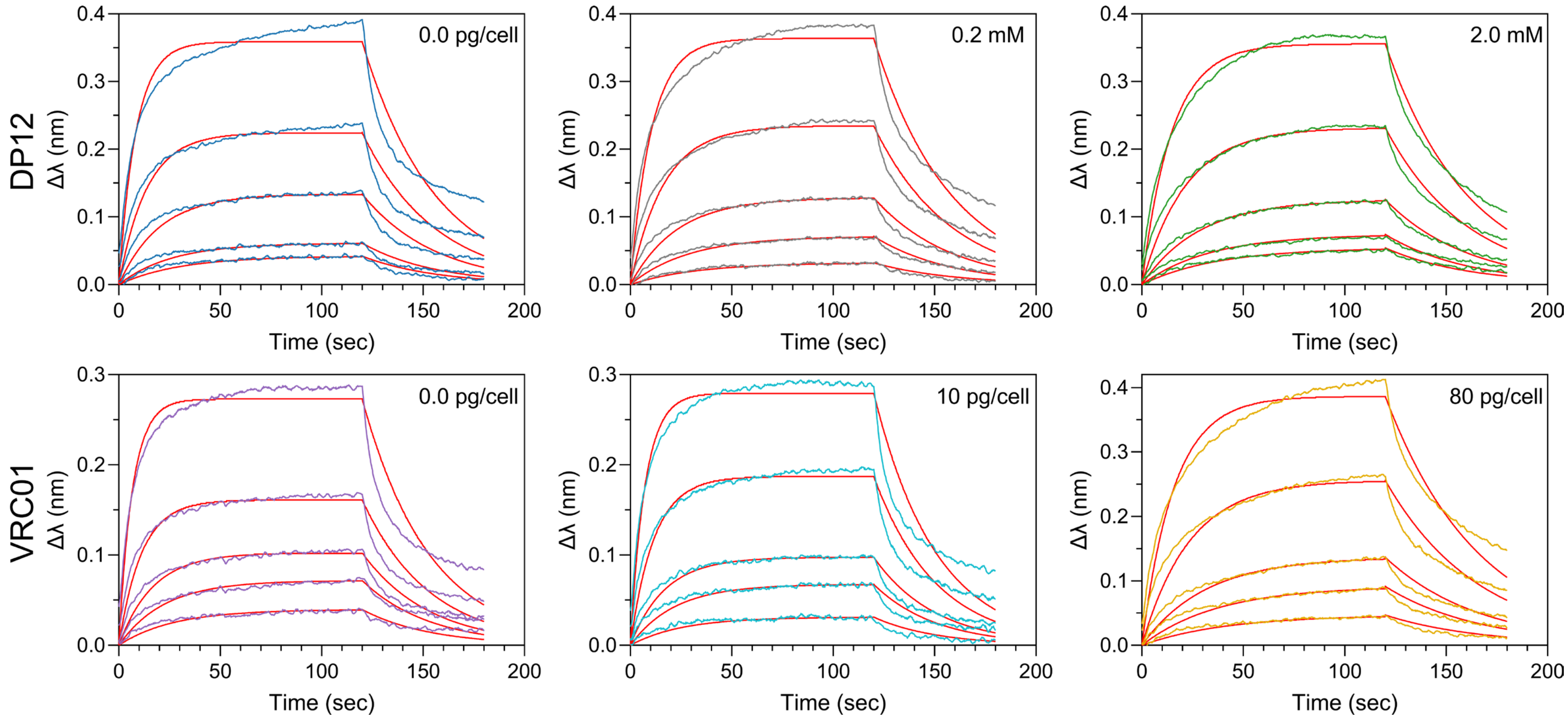


**Supplementary Figure 6. Bio-layer interferometry sensorgrams for CD16a:mAb binding kinetics assays**

Sensorgrams for mAb samples generated with different 2FG doses (indicated in the upper right-hand corner of each panel) for DP12 cells exposed to 10:90 split feeding (top row) and VRC01 cells fed with cell-normalised enhanced feeding (bottom row) are shown. The solid red lines correspond to the best fits to a 1:1 binding model using the Octet® Data Analysis software (version 12.0.1.2), while the coloured lines represent wavelength shifts for all assayed mAb concentrations.

| | $K_D$ | Aglyc. | HM | Fuc | Gal |
|---|---|---|---|---|---|
| $K_D$ | 1.00 | | | | |
| Aglyc. | 0.90 | 1.00 | | | |
| HM | 0.86 | 0.98 | 1.00 | | |
| Fuc | -0.87 | -0.99 | -1.00 | 1.00 | |
| Gal | -0.96 | -0.88 | -0.90 | 0.88 | 1.00 |

**Supplementary Figure 7. Correlation matrix across glycosylation features and CD16a:mAb binding affinity ($K_D$)**
